# Single-cell transfer RNA sequencing reveals principles of translational regulation

**DOI:** 10.64898/2026.11.11.765974

**Authors:** Laura C. Kida, Konstantin Helmsauer, Benan N. Nalbant, Austin A. Varela, Sydney B. Blattman, Yan-Ting Chen, Kaden Southard, Eli Metzner, Allison J. Vela, Meghana Pagadala, Rico C. Ardy, Hailey V. Goldberg, Meril Takizawa, Karolis Kumpaitis, Marisa Mariani, Catherine Snopkowski, Nabih Maslah, Santosha A. Vardhana, Susan DeWolf, Scott W. Lowe, Dana Pe’er, Ronan Chaligné, Leif S. Ludwig, Thomas M. Norman, Caleb A. Lareau

## Abstract

Transfer RNAs (tRNAs) decode genetic instructions from messenger RNA (mRNA) into protein, yet their abundance and regulation across cell states remain largely undefined, as multimodal measures of cellular tRNAs have remained elusive. Here, we develop a massively parallel assay for single-cell translatome and RNA sequencing (sctRNA-seq), which profiles transcriptome-wide mRNA and tRNA abundance simultaneously and chart the tRNA composition of 364,841 cells. Through multimodal perturbation screening of protein-coding genes and tRNA loci, we reveal dynamics across the mRNA and tRNA transcriptomes, delineating distinct tRNA-linked modules regulated by metabolic and Pol III-mediated transcription. Further, we uncover a close correspondence between codon demand and tRNA-mediated anticodon composition during hematopoietic multilineage commitment and differentiation. Tri-modal profiling of mRNA, tRNA, and proteins from primary human T cells reveals tRNA-mediated regulation that partially explains the discordance between mRNA and protein levels, particularly for selenoproteins and transcription factors. Finally, we uncover a stress-responsive tRNA module defined by gene-internal promoter sequences. Together, our analyses resolve tRNAs as a previously undefined layer of translational control underlying cellular identity and cell-state transitions.

## Introduction

Transfer RNAs (tRNAs) mediate the fundamental decoding of messenger RNAs (mRNAs) into proteins at the ribosome to complete the final step of the Central Dogma (**Fig. 1A**, (*1*, *2*)). While the human codon logic comprises only 61 principal codon species mapping to the canonical amino acids, the totality of cellular tRNAs is considerably more complex, with 432 genomic loci encoding tRNA molecules that additionally undergo post-transcriptional processing (*3*, *4*). As translating mRNA requires dozens to hundreds of tRNA molecules to decode a single peptide by design of the genetic code, the cellular tRNA pool exceeds an estimated 10^8^ copies in mammalian cells (*5*, *6*). Despite the remarkable abundance and foundational role tRNAs orchestrate in nearly all cells, cellular tRNA composition and regulation remain largely unexplored, particularly in cellular differentiation, disease, and perturbation.

**Figure 1.**
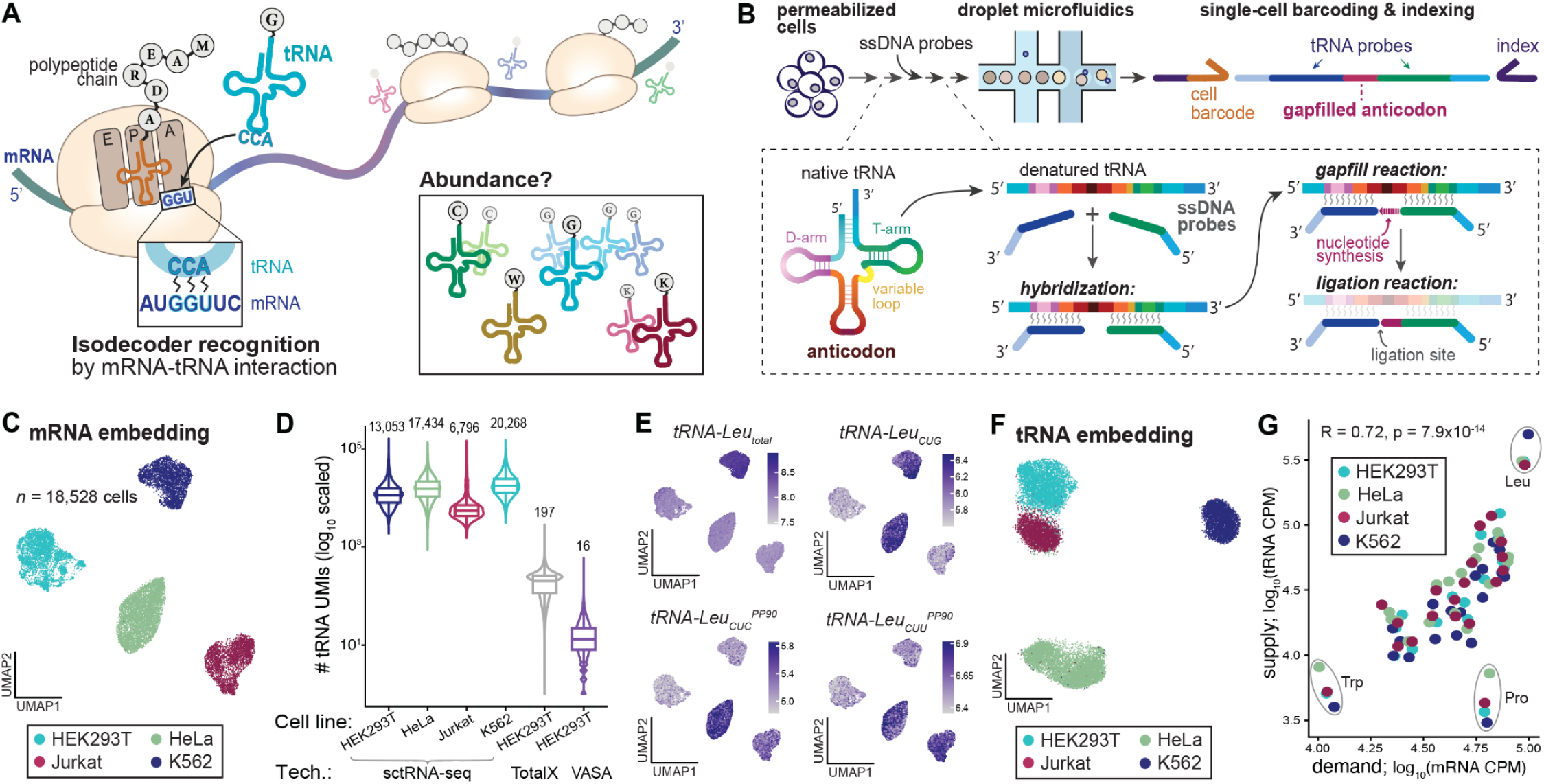
Design of sctRNA-seq. **(A)** Schematic of tRNA function during protein translation. **(B)** Schematic of sctRNA-seq, highlighting probe hybridization, gapfilling of the anticodon, and cellular barcoding via droplet microfluidics. **(C)** UMAP embedding of four cell lines annotated by cell line identity. **(D)** Quantification of tRNA detection across single-cell technologies. Labels: mean tRNA counts per cell per condition. **(E)** Annotation of variable tRNA species across different cell lines. **(F)** UMAP embedding of the same data as (C) but using only tRNA features. **(G)** Coordination of amino acid demand (based on mRNA codon counts) and supply (based on tRNA counts). Statistical test: Spearman correlation.

Recently, assays that quantify bulk cellular populations have characterized tRNA dynamics and functional variation in heterogeneous settings. Emerging work shows that the tRNA pool, much like corresponding mRNAs and proteins, is actively reprogrammed during cellular states, including cellular proliferation, coordination of pathogenic gene programs in disease (*7*), and translation of pro-metastatic gene programs during tumor evolution (*8*, *9*), including tRNA dosage effects (*10*). As one mechanism for this layer to contribute to the regulation of gene and protein expression, relative levels of tRNAs, such as changes in synonymous anticodons can dictate translational efficiency, fidelity, and protein folding (*11*).

While these anecdotes suggest that tRNAs act as a critical regulatory lever underlying cellular phenotypes, tRNA regulation has been largely understudied due to technical limitations in existing workflows for studying the biology of these RNAs. Unlike protein-coding genes transcribed by RNA polymerase II (Pol II), nuclear-encoded tRNAs are transcribed by RNA polymerase III (Pol III) through distinct machinery and regulatory mechanisms (*12*, *13*). The elements governing Pol III activity including tRNA gene-internal promoters known as A and B box present as conserved sequences across the whole tRNAome, yet the reported abundant shifts of transcript counts within the tRNA pool suggest an underlying regulatory logic. Because most single-cell RNA sequencing (scRNA-seq) technologies prime reverse transcription (RT) reactions from polyadenylated tails absent from tRNAs, tRNA profiling has largely been incompatible at single-cell resolution. While *in vitro* poly-A tailing methods enable non-zero yet low capture of tRNAs (*14–16*), comprehensive profiling of tRNA regulation and of the translatome–the coordinated dynamics of mRNA, tRNA, and proteins comprising the central dogma–has remained elusive.

Here, we introduce single-cell translatome and RNA sequencing (sctRNA-seq), which enables scalable co-quantification of mRNA and tRNA abundance by enhancing a widely adopted single-cell platform that can be further extended to detect surface and intracellular proteins, as well as genomic perturbations. In brief, through a computationally optimized design of single-stranded DNA (ssDNA) probes in combination with a short RT reaction, sctRNA-seq yields thousands of tRNA UMIs per cell with concomitant quantification of mRNA, proteins (via oligo-barcoded antibodies (*17*)), and guide RNAs (gRNAs) that collectively unveil the variation in the tRNA pool as an understudied layer of cell state regulation. Our workflow yields tRNA quantification with orders of magnitude greater sensitivity than approaches using functional polyadenylation or random priming (*14–16*, *18*), enabling a comprehensive assessment of the tRNA pool in thousands of single cells. We demonstrate the broad applicability of sctRNA-seq in decoding tRNA responses to targeted cellular perturbations (via Perturb-seq (*19*)) as well as differentiation and stimulation of primary human cells, revealing a model where tRNA pools are upregulated in response to increasing codon demands, and the cellular variability of tRNA pools partially explains the discordance between mRNA and protein abundances. Together, these vignettes demonstrate sctRNA-seq as an information-rich assay to uncover principles of translational regulation previously inaccessible in conventional single-cell profiling.

### ssDNA probes enable concomitant profiling of mRNAs and tRNAs

Despite their functional significance in decoding nucleic acids into functional peptides (**Fig. 1A**), sensitive quantification of tRNAs in single cells has been largely intractable. As standard RNA sequencing approaches require the RT of native RNAs into cDNA libraries, the extensive post-transcriptional modifications and internal base pairing of tRNAs collectively interrupt polymerase processivity (*20*). Recently, we (*21*, *22*) and others (*23*, *24*) have established orthogonal scRNA-seq assays that use thousands of ssDNA oligonucleotides to extend the 10X Genomics Flex kit (*25*), with several practical advantages. Instead of an RT step over the full RNA, the hybridization and barcoding of ssDNA pairs enables scalable profiling of Formalin-Fixed Paraffin-Embedded (FFPE) tissues (*25*), rare populations defined by transcriptomic features (*22*), and specific somatic mutations irrespective of the location on the transcript (*21*, *23*). Extending this logic, we conceived an assay to profile tRNAs with consecutive 25-nucleotide ssDNA probes that surround the tRNA anticodon (**Fig. 1B, Methods**). Using our recently characterized strand displacement deficient *Sulfolobus islandicus* polymerase (*21*), we leverage a high-fidelity gap-filling of the three anticodon bases between the pairs of ssDNA probes to infer base-pairing at the anticodon. These steps occur alongside the standard whole transcriptome amplification (WTA) panel, yielding concomitant profiles of cellular mRNAs and tRNAs (**Fig. 1B**). Collectively, sctRNA-seq comprises ssDNA hybridization, gap-filling, single-cell barcoding, and sequencing of concomitant mRNA and tRNA profiles (**Fig. 1B**).

To implement sctRNA-seq, we designed an optimized set of ssDNA probes to delineate the cellular tRNA pool. We implemented a greedy algorithm to maximize the number of unique probe pairs that could hybridize to cellular tRNAs while accounting for both redundancy in genomic tRNAs and mismatch tolerance in the ssDNA probes (**Fig. S1A, Methods**). Using the set of all 432 canonical tRNA loci in the human genome (*4*), our algorithm identified 145 unique ssDNA probes that would hybridize on the 5’ end of the anticodon and 89 on the 3’ end of the anticodon, both achieving significantly higher sequence coverage than alternative approaches, yielding 181 high-confidence distinct probe pairs, 21 more than without the greedy heuristic (**Fig. S1B,C,** permutation test *p* = 4.5×10^-21^**, Methods**). Coupled with the high-fidelity 3-nucleotide RT step over the anticodon, our approach yields 217 combinatorial canonical features reflecting the diversity of the cellular tRNA pool, which maps the breadth of possible cellular tRNA decoding (**Fig. S1D, Methods; Supplemental Table 1**). Notably, the accuracy of the *Sulfolobus* polymerase allows us to confidently differentiate all possible isodecoders, including the 13 arising from post-transcriptional modifications (*21*). We appended our computationally optimal probe panel for decoding tRNA variability to the commercial whole transcriptome amplification (WTA) panel for mRNA profiling in all subsequent experiments.

We established this assay using a mixture of four human cell lines: HEK293T, HeLa, Jurkat, and K562 (**Fig. 1C**). Similar to genotyping mutations on RNA molecules (*21*), we found that including betaine in the gap-filling step substantially increased the diversity of tRNA molecules profiled, including representation of non-genetically encoded anticodons and mature tRNAs with excised introns (**Fig. S1E-G**). From these profiles, we confirmed the fidelity of our gap-filling reaction and overall assay design, as 99% of sequenced tRNA molecules in all cell lines contained a resulting three-nucleotide gap-fill product (**Fig. S1I**). Collectively, these results confirm the sensitivity and accuracy of our assay using computationally optimal ssDNA probes that hybridize to tRNA molecules.

We confirmed known mRNA expression discriminated each cell line, including *CD3E*, *CXCR4*, *VIM*, and *BCR* (**Fig. S1H**). Compared to other single-cell tRNA profiling methods, sctRNA-seq recovers ∼70-900-fold more tRNA UMIs per cell across tens of thousands of cells (**Fig. 1D**). This gain in throughput and resolution reveals hierarchical differences in tRNA expression at four levels: total tRNA species abundance, anticodons, isodecoders, and mature tRNA molecules with post-transcriptional modifications at the wobble position for leucine and isoleucine as exemplar species (**Fig. 1E**, **Fig. S1K**). Dimensionality reduction, clustering, and two-dimensional projections using probe-level tRNA features similarly separated cell lines as a canonical mRNA embedding (**Fig. 1F**). Overall tRNA availability per amino acid closely correlated with cellular demand from mRNAs (*r* = 0.72, *p* = 7.9×10^-14^, **Fig. 1G, Methods**). Proline tRNA abundance was an outlier in all four lines, consistent with previous reports of slow proline incorporation at the ribosome relative to other tRNAs (*26*). While each tRNA species was well-represented in the various lines, we reasoned that the overall tRNA levels in a cell may serve as a proxy for overall translational activity, reflecting lower activity in Jurkats than K562s, which we confirmed through puromycin incorporation and flow cytometry via SCENITH (**Fig. S1J; Methods**) (*27*). For individual tRNA genes, pseudobulk expression matched Pol III abundance as measured using ChIP-seq and at the anticodon level, it was comparable to orthogonal RT-based bulk tRNA-seq libraries, broadly corroborating the accuracy of our tRNA quantification (**Fig. S1L-N; Methods**). Overall, the unique design of sctRNA-seq enables sensitive and accurate quantification of individual cellular tRNA profiles through a facile extension of a widely used commercial assay.

### Validation of sctRNA-seq by parallel perturbation

As sctRNA-seq extends the Flex assay, where concomitant detection of gRNAs has been described (*28*, *29*), we reasoned that parallel perturbation of tRNA loci could be achieved with concomitant profiling of mRNAs and tRNAs alongside guide RNA (gRNA) detection, enabling an assay benchmark (**Fig. 2A**). We selected exemplar genomic loci that transcribe tRNAs encoding different amino acids, including selenocysteine (SeC) and initiator methionine (iMet) as well as the canonical 20 amino acids. We designed a total of 73 guides, including 3 for most loci and 10 non-targeting guides (**Table S2; Methods**).

**Figure 2.**
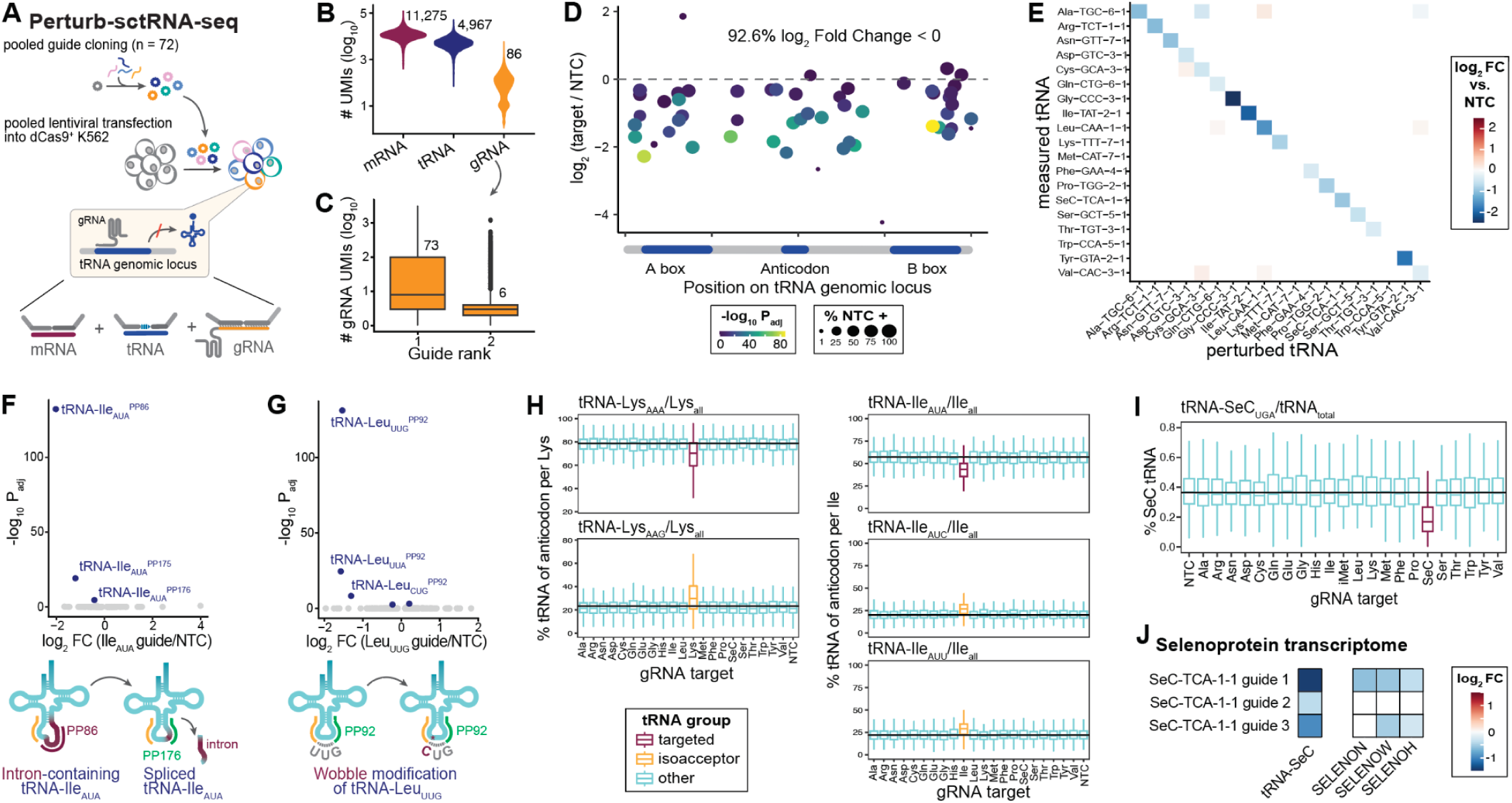
Sensitivity and specificity of sctRNA-seq assessed via targeted perturbations. **(A)** Schematic for CRISPRi-based validation of sctRNA-seq, including pooled cloning and profiling of mRNA, tRNA, and gRNAs in the same cells. **(B)** UMI abundance of mRNAs, gRNAs, and tRNAs across the whole cell population (n = 10,899 cells). Numbers reflect median values. **(C)** UMI abundance of top gRNAs, verifying high-quality assignments. Numbers reflect median values. **(D)** Effect of perturbation relative to the position of the gRNA target. Bottom: Relative positions of landmark tRNA features. **(E)** Effect of gRNA perturbation on all targeted tRNAs with unique probe-mapping. **(F)** Differential tRNA effects for Ile_AUA_ perturbation, noting three probe pairs reflecting intron-containing and spliced tRNA species. **(G)** Differential tRNA effects for Leu_UUG_ perturbation, noting a probe pair with germline-encoded and post-transcriptional (wobble) modifications. **(H)** Effect of tRNA perturbation altering specific isodecoders for Lysine (left) and Isoleucine (right). Box plot displays the fractional abundance of each anticodon within each isoacceptor family (% isodecoder per isoacceptor), colored by whether the respective anticodon is the guide’s on-target isodecoder (red), on-target isoacceptor but off-target isodecoder (yellow) or not (other, blue). **(I)** Effect of selenocysteine (SeC) tRNA perturbation as a proportion of SeC tRNA relative to all anticodons. **(J)** Effect of tRNA-SeC perturbation on selected mRNAs encoding selenoproteins. Boxplots: center line, median; box limits, first and third quartiles; whiskers, 1.5× interquartile range.

Pooled transfection and profiling of 10,899 cells yielded high-quality data metrics per modality, including a mean 5,615 tRNA UMIs per cell (**Fig. 2B**) and clear perturbation-assignment of highest-scoring guide identities by UMI counts per cell (**Fig. 2C**). Instead of conventional CRISPRi guide designs for protein-coding genes that disrupt the promoter upstream of the target gene, we designed gRNAs targeting tRNA genes intragenically at tRNA-associated Internal Control Regions (ICRs) and found the guide effect to be independent of position across the tRNA loci (**Fig. 2D**). Encouragingly, we observed at least a two-fold knockdown for 84.2% of loci as well as gene-level specificity of individual tRNAs (**Fig. 2E, S2A**). These results confirmed that both Pol III-encoded transcriptional products could be silenced by the widely-used dCas9-ZIM3 epigenome editor and validates sctRNA-seq as a sensitive readout for primary tRNA perturbation.

A feature of our assay design included the ability to detect tRNAs across different stages of their life-cycle, including pre- (tRNA-Ile ^PP176^) and post- (tRNA-Ile ^PP86^) spliced forms of an isoleucine tRNA as well as the germline (tRNA-Leu ^PP92^) and post-transcriptionally modified (tRNA-Leu ^PP92^) forms of leucine, all of which responded to CRISPRi perturbation (**Fig. 2F,G**). Remarkably, despite the genomic redundancy of tRNA-encoded loci, we found that single guide perturbations are detectable through the overall composition of the cellular tRNA pool, as the targeted species decreased by 5-10% within isodecoder families with accompanying increases of other isodecoders (**Fig. 2H**, **Fig. S2B,C; Methods**). An exception to tRNA germline redundancy is the selenocysteine tRNA (tRNA-SeC), which is encoded at a single locus on chromosome 19. Across the three guides designed to target the tRNA-SeC locus, we observed a 44.7% knockdown of SeC as a proportion of the cellular tRNA pool (*p* = 1.7×10^-47^) with a clear dose-response in guide efficacy of both the primary (i.e., tRNA transcription) and secondary readouts (i.e., abundance of mRNA encoding selenoproteins; **Fig. 2I,J**). Together, these parallel perturbations corroborate the accuracy and sensitivity of our assay for decoding changes in the cellular tRNA pool via sctRNA-seq.

### tRNA response to protein-coding perturbations

Despite nearly a decade of single-cell mRNA and gRNA profiling via Perturb-seq (*30*, *31*), the regulatory factors of protein-coding genes that impact Pol III-mediated tRNA transcription remain largely unexplored. To characterize the joint effects of coding-gene perturbations on mRNA and tRNA regulation, we selected 174 guides targeting genes across Pol II/III transcriptional machinery, metabolic regulatory pathways (i.e. MTOR) as well as tRNA modification and maturation (i.e. tRNA synthetases) from a genome-wide Perturb-seq screen (*31*) with an additional 68 guides targeting chromatin regulatory machinery (**Fig. 3A; Methods; Table S3**). We transfected a lentiviral pool of these 242 targeting guides and 18 NTCs into dCas9-ZIM3 K562s, yielding 89,102 cells with high-quality mRNA, tRNA, and gRNA capture and validated the anticipated on-target gRNA activity (**Fig. S3A, Methods**). Pairwise correlation of mRNA data from sctRNA-seq recovered previously established clusters of target gene perturbations, including pathways related to tRNA charging, tRNA modifications, and cell regulation via mTOR (**Fig. 3B, Methods**). Clustering tRNA responses according to perturbation recovered a similar overall structure but with additional co-variation linking components of the Pol III machinery, including components of the general transcription factor 3 (*GTF3*) family, *BDP1* (a subunit of GTF3B), and *TRMT5* (a tRNA methyltransferase)(**Fig. 3C**). Differential expression analyses of the tRNA and mRNA modalities interestingly showed largely uncorrelated results, where Pol III-associated genes dynamically regulated nearly 100 tRNA species but comparably few mRNAs (**Fig. 3D**). Conversely, genes previously characterized to trigger the integrated stress response (ISR), including *HSPE1, QARS, and AARS*, resulted in hundreds of differentially expressed mRNAs with corresponding tRNAs also changing in response to these perturbations. For perturbations that strongly affected tRNAs, most resulted in an overall decrease in tRNA expression, except *TRMT5*, which catalyzes a methyl transfer reaction specifically when guanine is in position 37 of the tRNA, as reflected in a tRNA-specific reaction (**Fig. 3E; Fig. S3B-D**).

**Figure 3.**
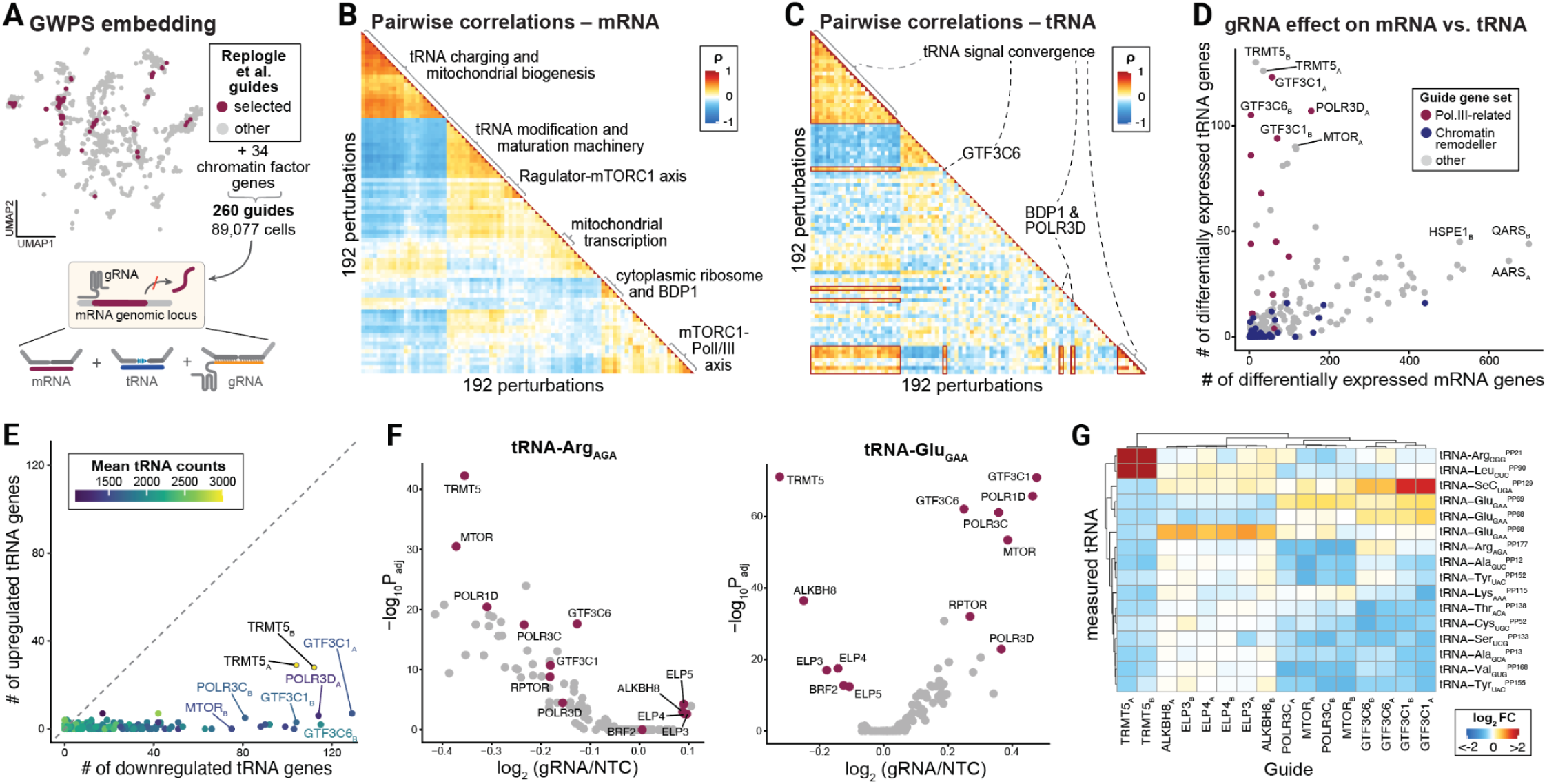
Epigenetic perturbation of protein-coding genes results in coordinated tRNA changes. **(A)** Reduced dimensionality of GWPS conditions. Selected genes for the perturbation experiment are noted. **(B)** Pairwise transcriptomic correlation of all 260 guides. Selected functional guide clusters are labeled. Colored by z-score of fold changes compared to NTC guides (ρ, Spearman correlation). **(C)** Pairwise correlations of guide perturbation profiles based on probe-level tRNA counts. Same row and column order as in B. **(D)** Enumeration of differential mRNAs and tRNAs per individual gRNA target. **(E)** Correlation plot of the number of down- (x-axis) and up-regulated (y-axis) tRNA gene products that are differentially regulated by each guide target, colored by the mean total cellular tRNA count per guide. **(F)** Volcano plots of tRNA-Arg_AGA_ (left) and Glu_GAA_ (right) changes in response to different noted perturbations. **(G)** Summary of perturbation-specific effects on tRNAs for selected guides relative to NTCs.

Beyond these global changes in tRNA expression, we observed perturbation-induced changes in specific tRNAs that were consistent across members of other protein complexes. These included concordant isodecoder differential expression; abundance compared to NTC was consistent across contextual members of protein complexes such as the tRNA transcriptional machinery; and the elongator complex proteins (ELP1-3) responsible for catalyzing modifications in the wobble position of certain tRNAs with isodecoder-specific directionality (**Fig. 3F**). Clustering among isotranscripts recovered the magnitude of these tRNA perturbations across substrates and non-substrates for the GTF gene group and tRNA-modifying enzymes such as TRMT5 and ALKBH8 (**Fig. 3G**). While perturbing chromatin accessibility factors showed negligible effects on the tRNA pool, knockdown of cytosolic and mitochondrial tRNA synthetases produced a clear signature that recovered the compartmental identity of the guide target (**Fig. S3E,F**). Notably, cytosolic synthetase perturbations strongly altered tRNA levels, yet the isoacceptor-level signature was largely conserved across both groups rather than reflecting the cognate synthetase substrates. This suggests a general stress response to synthetase perturbation, not an isoacceptor-specific effect. Together, these analyses demonstrate the utility of our platform for scalable perturbation analyses to uncover previously underappreciated regulatory principles underlying tRNA expression.

### Co-transcriptional dynamics of the transcriptome and tRNAome in primary cell differentiation

As multilineage cellular differentiation results in thousands of changes to the cellular transcriptome and proteome, we hypothesized that cellular tRNA composition is similarly reprogrammed to support these distinct cellular modules. We thus applied sctRNA-seq to an *in vitro* differentiation model of multi-lineage hematopoietic differentiation with continuous trajectories from a hematopoietic stem and progenitor cell (HSPC) population to three distinct lineages. We expanded and differentiated human CD34^+^ HSPCs using cytokine cocktails and profiled 71,922 cells, yielding populations that expressed canonical marker genes in the erythroid (*GATA1*), monocytic (*CD14*), and megakaryocytic (*PF4*) lineages (**Fig. 4A-C; Methods**). Strikingly, we observed similar lineage-specific expression of individual tRNA isotranscripts akin to these canonical marker genes, including tRNA-Val_GUC_^PP163^ (erythroid), tRNA-Cys_UGC_^PP56^ (monocytic), and tRNA-Ile_AUC_^PP82^ (megakaryocytic; **Fig. 4D**). In addition to these marker gene-like properties, we observed a decline in tRNA abundance over HSPC differentiation as well as differences between the same amino acid-containing but functionally divergent methionine isodecoder tRNAs tRNA-Met_AUG_^PP120^ and tRNA-iMet_AUG_^PP172^ as well as closely related tRNA species such as tRNA-Asp_GAC_^PP44^ and tRNA-Asp_GAC_^PP45^ (**Fig. S4A-C**).

**Figure 4.**
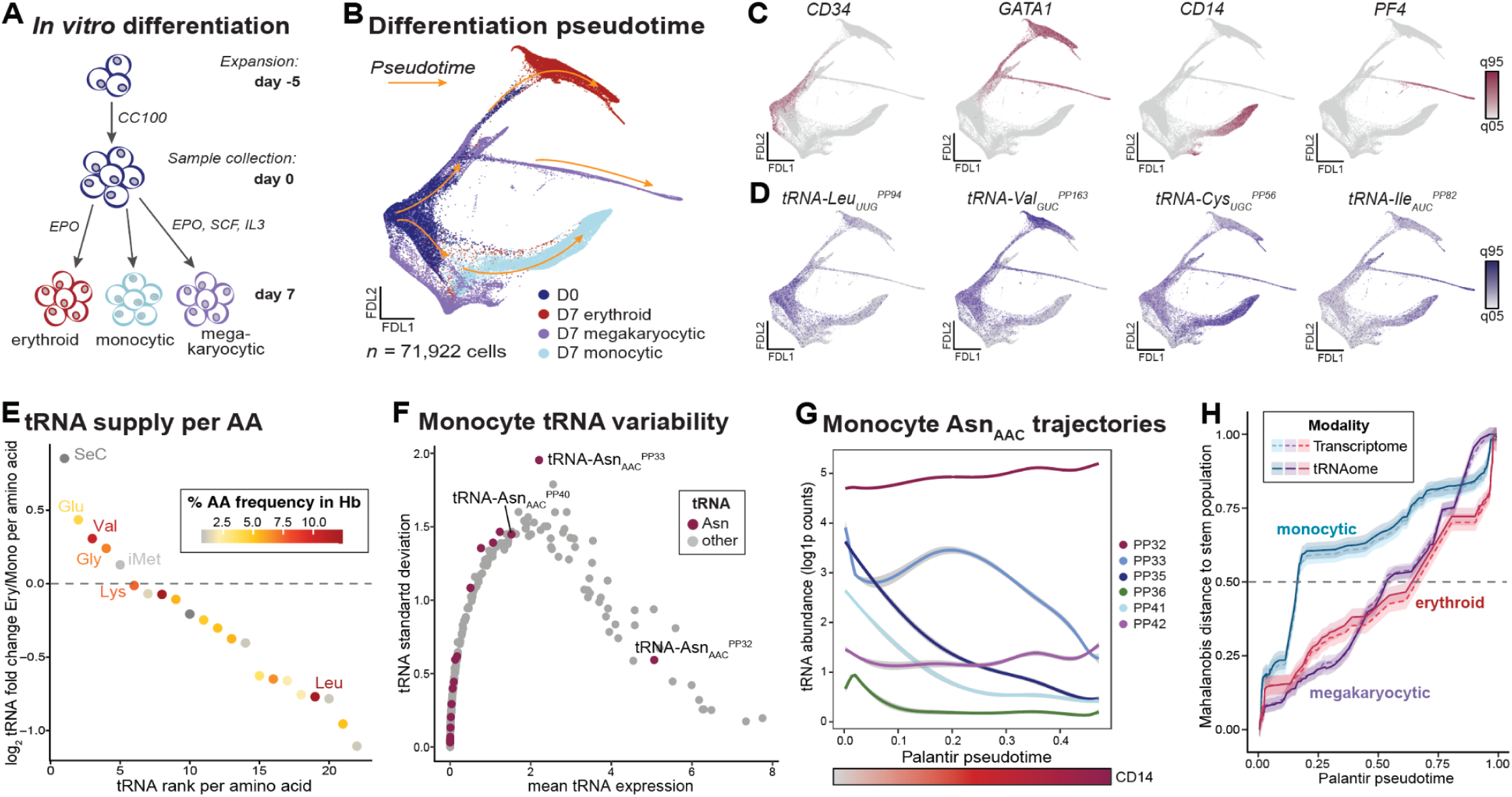
Continuous regulation of tRNA abundance mirrors mRNA changes over multi-lineage differentiation. **(A)** Schematic of *in vitro* differentiation of CD34^+^ selected primary cells. **(B)** Palantir pseudotime plotted on a force-directed layout (FDL). Arrows indicate the direction of the pseudotime differentiation trajectory. **(C)** Lineage-specific transcriptomic markers for stem and progenitor cells as well as erythroid, monocytic, and megakaryocytic lineages. **(D)** Same as (C) but for lineage-enriched tRNAs. **(E)** Amino acid-level tRNA supply between differentiating erythrocytes and monocytes, colored by the prevalence of each amino acid in human hemoglobin. **(F)** Expression and standard deviation of individual tRNA features in differentiating myeloid cells, highlighting Asn_AAC_ tRNAs. **(G)** Trajectories of tRNA-Asn_AAC_ isodecoders over pseudotime. **(H)** Mahalanobis distances across pseudotimes of mRNA and tRNA profiles relative to the stem and progenitor populations. Color shades represent 95% confidence intervals.

We reasoned that the highly regulated cell-type specificity of the proteome would dictate many of these dramatic changes observed in the composition of the tRNA landscape. Indeed, we observed a two-fold increase in selenocysteine tRNAs in committed erythroid progenitors relative to monocytic progenitors, potentially reflecting the synthesis demand of SeC-containing proteins including thioredoxin reductases and glutathione peroxidases to protect against reactive oxygen species (ROS) (*32*, *33*) (**Fig. 4E**). We further observed an upregulation of glycine and valine-decoding tRNAs in erythroid cells, which may reflect their relative enrichment of the erythroid proteome, including the specialization towards the synthesis of **α**- and **β**-globin (*34*).

We next sought to characterize the compositional evolution of these lineage-specific tRNA pools along the differentiation trajectory. Based on our data and previous reports on tRNA dynamics in differentiation (*35*, *36*), we reasoned that these anecdotal and systematic changes in the tRNAome may reflect a continuous differentiation reprogramming of the tRNA pools reminiscent of other specific molecules, including accessible chromatin (*37*), mRNA (*38*), and surface proteins (*39*). For each differentiation trajectory, we identified the most variable tRNAs, including pronounced regulation of the asparagine-decoding tRNAs (**Fig. 4F**), and computed the variability of individual isodecoders across pseudotime, noting pronounced divergence of tRNA-Asn_AAC_ isotranscripts across the monocytic cell population, while other lineages featured other variable tRNAs (**Fig. S4D**). Among tRNA-Asn_AAC_ isotranscripts, we observed locus-specific dynamics with variable slopes, suggesting a higher-order regulatory process beyond simple transcriptome-derived codon demand (**Fig. 4G**).

Across all three lineages, we found matched trajectories of tRNA and lineage-defining mRNA genes (**Fig. S4E-G**). Motivated by evidence of lineage priming from epigenomic profiling (*40*), we sought to determine whether the large codon-demand changes during differentiation preceded, followed, or were coordinated with the anticodon supply provided by tRNAs. Using codon demand (via mRNA) and supply (via tRNA), we computed compositional distances along each differentiation trajectory relative to the multi-lineage stem and progenitor cells (**Methods**). Across all three lineages, we observed that tRNAome remodeling over pseudotime tightly coincided with changing codon demands determined by protein-coding genes, an insight robust to data permutation and hyperparameter changes (**Fig. 4H; Fig. S4H; Methods**). These data support a model where transcriptional regulation during differentiation coordinates both Pol II and Pol III-regulated transcripts to meet dynamic protein synthesis requirements in fate commitment.

### Regulation of tRNAs mediates protein abundance

The discordance between mRNA and protein abundance is widely observed in many systems (*41–43*). As tRNAs are the central decoders between genetic instructions and observed protein levels, we reasoned that variation among cellular tRNA pools may underlie an axis of this variation. With the inherent modularity of our assay, we extended sctRNA-seq using a panel of 344 oligo-conjugated antibodies (**Table S4**) for simultaneous quantification of mRNA, tRNA, and protein abundance in the same single cells, profiling 52,798 primary T cells at varying stages of T cell receptor stimulation (**Fig. 5A,B, Methods**). Dimensionality reduction and clustering showed an expected split of the cell population according to experimental stimulation condition (**Fig. 5C**). Our proteomic panel recapitulated previously annotated proteomic changes in these conditions, including an upregulation of exhaustion-related markers, such as PD1 and CD155, in the chronically stimulated T cell condition relative to the acute stimulation, which in turn had higher expression of markers of activation, (i.e., CD80 (B7-1), CD86 (B7-2), and CD56, **Fig. S5A**). Reminiscent of lineage-specific tRNAs emerging during hematopoietic differentiation, we similarly observed tRNA changes within these distinct T cell populations, further reflecting previously underannotated tRNA changes in variable cell states (**Fig. S5B**). We next quantified the overall per-cell ratio of tRNA to mRNA, observing this ratio to increase after stimulation (**Fig. 5D**). Correspondingly, counts for tRNA-iMet were elevated in the acutely stimulated D8 T cell sample (D8a/D2: log_2_FC = 0.7, D8a/D8c: log_2_FC = 1.1), likely reflecting higher overall translational activity in these cells due to the central role of tRNA-iMet in the initiator complex (*44*) and previously reported increased cell proliferation in tRNA-iMet overexpression (*45*, *46*)(**Fig. 5D, E**).

**Figure 5.**
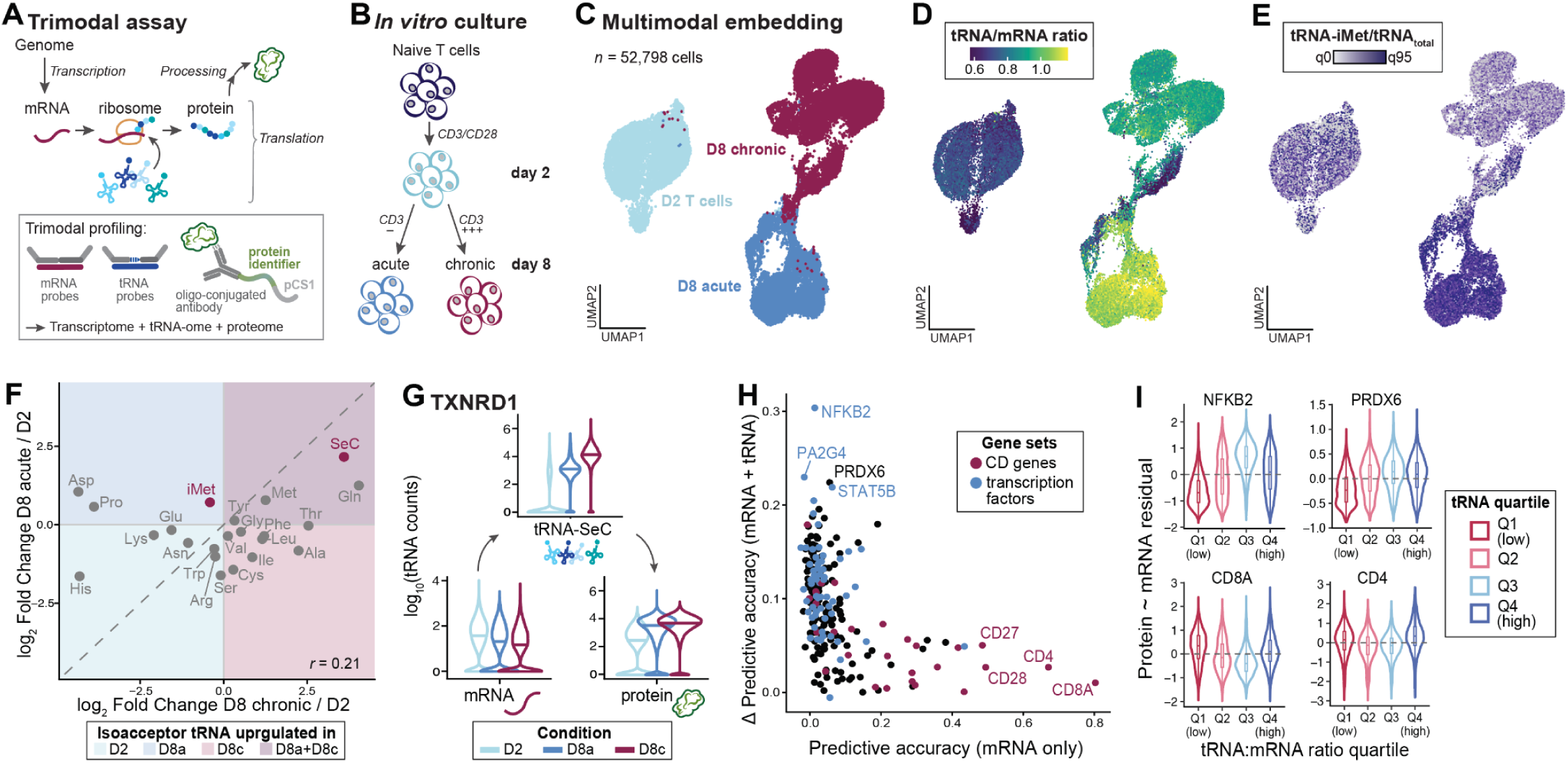
Variable tRNA levels regulate protein abundance in single cells. **(A)** Schematic of the extended sctRNA-seq assay into a trimodal readout including protein quantification via oligo-conjugated antibodies. **(B)** Schematic of *in vitro* setting for T cell stimulation. Naive T cells were activated with CD3/CD28 and profiled at day 2. They were then subjected to acute (CD3+) or chronic (CD3+++) stimulation and profiled at day 8. **(C)** UMAP of the weighted-nearest-neighbor multimodal embedding (n = 52,798 cells), coloured by condition: D2, D8 acute (D8a), D8 chronic (D8c). **(D)** Per-cell ratio of total tRNA to mRNA counts. **(E)** Fraction of initiator tRNA-iMet among total tRNAs detected per cell. **(F)** Isoacceptor-level tRNA fold changes relative to D2 under acute (y-axis) versus chronic (x-axis) stimulation. Highlighted are isoacceptor tRNAs iMet and SeC. **(G)** Distributions of *TXNRD1* mRNA (bottom left), tRNA-SeC (top), and TXNRD1 protein (bottom right) across conditions. The horizontal bar marks the median. **(H)** Correlation plot of the accuracy of predicting protein abundance from mRNA alone (x-axis) versus the gain in accuracy from adding tRNA features (y-axis), per protein. Coloured: Cell surface (CD) genes (red) and transcription factors (blue). **(I)** Residuals of protein abundance after regression on mRNA, stratified by quartile of the tRNA:mRNA ratio (Q1, lowest; Q4, highest).

Among genes with co-quantified mRNA and protein levels, we observed an apparent mismatch of mRNA- and protein-level abundance for markers including MYC or NFKB2 (**Fig. S5C**). Indeed, even within each stimulation group, low to negative correlations between modalities were apparent for both MYC (Spearman *r* -0.29-0.18) and NFKB2 (Spearman *r* 0.05-0.15)(**Fig. S5D**). We reasoned that the overall tRNA pool may help explain the apparent discordance between these measurements. To characterize the impact of tRNAs regulating protein abundance, we first assessed whether an individual tRNA species was particularly responsive to stimulation and could be critical in this system. We noted that tRNA-SeC was most strongly overexpressed across isoacceptors in both day 8 conditions compared to D2 (D8a: log_2_FC = 2.2, D8c: log_2_FC = 3.6) and particularly abundant in the chronically stimulated cells (**Fig. 5F, Fig. S5E**). Based on our characterization of regulated selenoproteins in our Perturb-seq experiment (**Fig. 2**), we reasoned that regulation of this tRNA may similarly affect selenoprotein transcript and protein expression in this primary cell system. Indeed, for the selenoproteins assayed in our protein panel and mRNA probe panel, we observed translational regulation in which increased tRNA-SeC explained increased protein production in chronically stimulated cells despite relatively lower mRNA levels for TXNRD1 (**Fig. 5G**) and GPX4 (**Fig. S5F**). This result suggests that T cells upregulate the tRNA-SeC transcript to produce selenoproteins to mitigate the ill effects of chronic stimulation as previously observed under tRNA-SeC knockout (*47*).

We next sought to generalize this observation for mRNA/protein pairs with less obvious tRNA bottlenecks, reasoning that a combination of isodecoders could influence overall translational efficiency. To assess this systematically, we computed the predictive accuracy of estimating protein expression from mRNA alone and from a combined model with mRNA and tRNAs, while adjusting for relevant covariates, including experimental condition and per-cell UMI counts for each respective modality (**Methods**). Then, for each gene, we fit a regularized linear model (lasso), with and without isodecoder-level tRNA measurements, to identify proteins regulated by variation in tRNA abundance. For each gene, we compared baseline accuracy from mRNA abundance with the added gain from the tRNA modality (**Fig. 5H**). This analysis yielded two notable protein populations with differing predictions. First, surface and lineage-specific markers such as CD8A, CD4, and CD28 showed stable, persistent protein expression that mRNAs modeled well. Conversely, transcriptional regulators such as NFKB2, STAT5B, and PA2G4 were largely uncorrelated with baseline mRNA but modeled more effectively when tRNA variability was included at the individual-cell level. These dynamics were partially recapitulated, as the residuals of the baseline model correlated with tRNA/mRNA ratios for genes better modeled by tRNAs (i.e., transcription factors) than by surface markers (**Fig. 5I, Fig. S5G**). Together, these analyses support a concept in which lineage-defining features may be more directly translated from underlying mRNA levels, whereas transcriptional regulators of dynamic cellular processes may be tuned by tRNA pools.

### Sequence-intrinsic tRNA responses under stress conditions

Because translational adaptation is a hallmark of cellular stress adaptation (*48*), we reasoned that combining CRISPR and chemical perturbations could resolve the dynamics of the cellular stress response. To achieve this, we applied the same panel of 260 guides to K562s cultured in amino acid-depleted (AAd) media or treated with mTORC1-inhibitor Torin-1 (**Fig. 6A; Methods**). We adapted sctRNA-seq for use with the droplet-based, combinatorial indexing Flex v2 kit to profile 103,372 cells across all three treatment conditions, yielding a mean 5,392 mRNA UMIs and 1,137 tRNA UMIs per cell. Treated cells showed the expected transcriptomic profiles, including upregulation of amino acid transporters and tRNA synthetases under starved conditions, whereas Torin-1-treated cells showed an upregulation of mitochondrial genes (**Fig. S6A**). This design allowed us to distinguish treatment-associated changes in tRNA profiles from the effects of individual genetic perturbations within each treatment condition (**Fig. 6B**).

**Figure 6.**
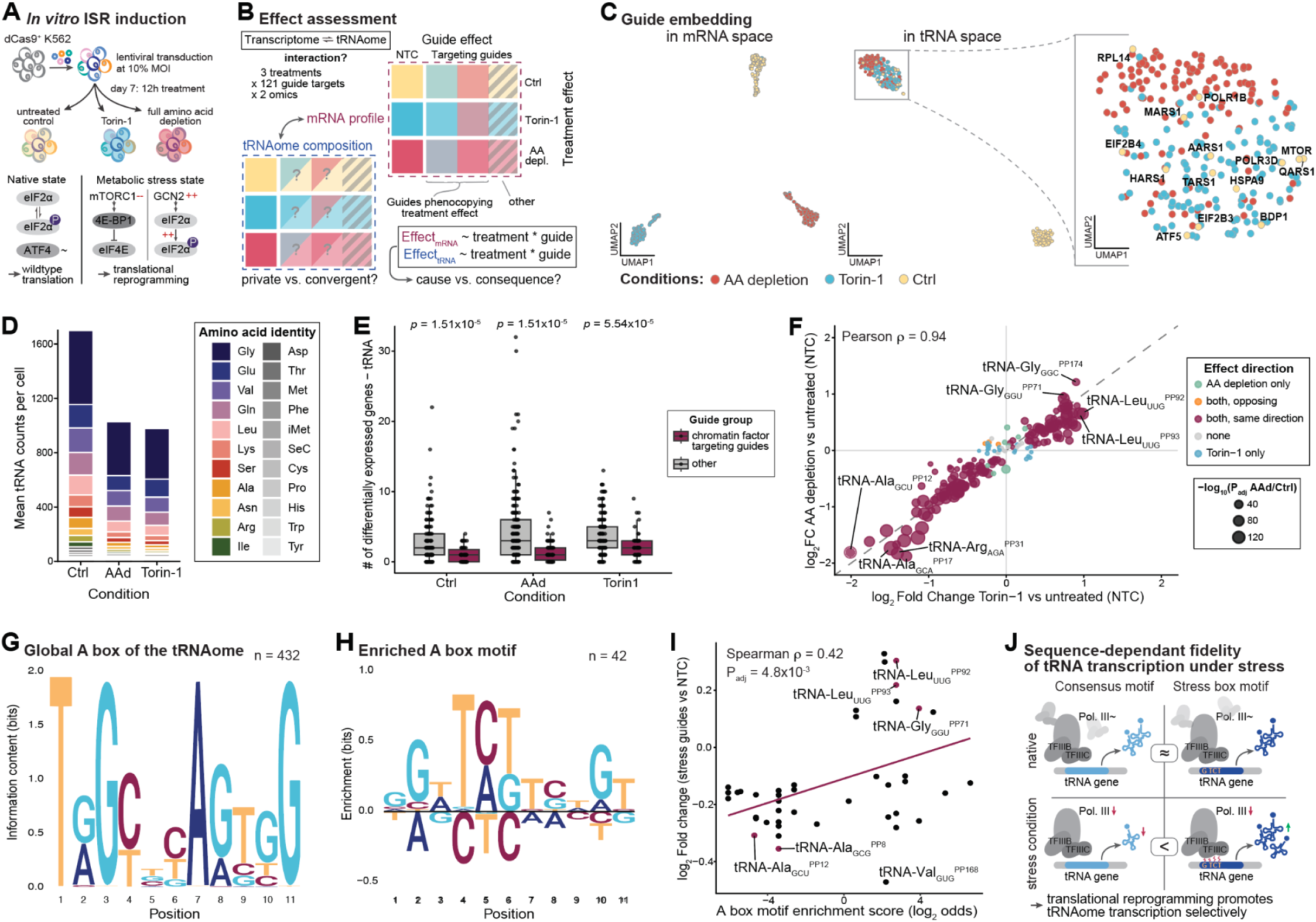
Sequence intrinsic features underlie stress-responsive tRNA transcriptional regulation. **(A)** Schematic of experimental design. K562 cells expressing dCas9 were transduced with a lentiviral CRISPRi library targeting 121 genes at ∼0.1 MOI. On day 7, cells were cultured for 12 hours under standard conditions (untreated control), with Torin-1, or in amino acid-depleted medium to induce cellular stress pathways. **(B)** Analytical framework of perturbation effects based on quantification of 3 treatments × 121 guide targets × 2 modalities (transcriptome and tRNAome). **(C)** UMAP embeddings of pseudobulked guide-level profiles in mRNA space (left) and tRNA space (right), colored by condition. Inset: magnified view of the tRNA-space stress cluster, with selected untreated perturbation targets labeled. **(D)** Mean tRNA counts per cell in non-targeting control (NTC) cells, stratified by amino acid identity. **(E)** Boxplot for the number of differentially regulated tRNA genes upon perturbation through chromatin factor-targeting (red) or other (grey) guides across treatment conditions. **(F)** Per-isodecoder log₂ fold changes in NTC cells comparing amino acid depletion (y-axis) against Torin-1 (x-axis). Point size indicates −log₁₀ adjusted P value for amino acid depletion vs. control. Pearson ρ = 0.94. **(G)** Sequence motif for the A box across the whole tRNAome. Letter height indicates information content (bits). **(H)** Enriched sequence motifs of the A box internal promoter elements of the 42 tRNA isotranscripts significantly upregulated in (F). Letter height indicates enrichment (bits). **(I)** Correlation between the A-box motif enrichment score and the enrichment motif from (H). **(J)** Schematic for the context-dependent relevance and quantitative consequences of different A box motifs for tRNA transcription.

To compare these responses across conditions and perturbations, we aggregated single-cell profiles into 393 metacells defined by treatment condition and guide target identity. This dual-perturbation setting produced distinct guide embeddings in mRNA and tRNA space. While transcriptomic signatures differed markedly between all three samples (Torin-1, AA depletion, untreated control), we observed a convergence of tRNA metacell phenotypes in this high-dimensional space (**Fig. 6C**). Here, untreated metacells separated from a cluster encompassing both stress-induced conditions. Notably, however, a subset of guides among untreated cells induced similar patterns of tRNA changes as seen across stress-induced conditions. Those guides targeted genes closely tied to the ISR, including *ATF5* and *EIF2B4* (*49*). Stress induction most prominently led to a significant decline in per-cell total tRNA counts (log_2_FC(AAd/control) = -0.70, log_2_FC(Torin-1/control) = -0.74, **Fig. 6D**). While the composition of the tRNA pool remained relatively stable among isoacceptor tRNA families (AAd vs. control: ρ = 0.99, *p* = 2.8×10^-6^; Torin-1 vs. control: ρ = 0.98, *p* = 3.3×10^-6^; **Fig. S6D,E**), we observed coordinated changes among isodecoder tRNAs across both treatments compared to the untreated control (ρ = 0.94, **Fig. 6F**). Correlating the tRNA changes on isotranscript-level, we observed a strong correlation of both treatments compared to untreated NTCs (Pearson ρ = 0.94), revealing tRNAs belonging to amino acids overall abundant in the human transcriptome including leucine (tRNA-Leu_UUG_^PP92^ and tRNA-Leu_UUG_^PP93^) and glycine (tRNA-Gly_GGC_^PP174^ and tRNA-Gly_GGU_^PP171^) encoding tRNAs (**Fig. 6F**).

We hypothesized that perturbations of chromatin-modifying genes would yield more apparent changes in tRNAs under these stress conditions. However, as in our initial protein-coding gene-targeting screen, knockdown of these genes did not produce systematic differential changes across the tRNA pool, affecting few tRNA genes overall, with stronger effects on the mRNA transcriptome (**Fig. 6E, Fig. S6C**). Because canonical chromatin remodeling agents had minimal impact on tRNA transcription, we sought to identify other mechanisms that could explain this stress-responsive tRNA signature. Perturbing tRNA-modifying factors we noted in our earlier screen (**Fig. 3**) affected tRNA abundance and revealed similar profiles under treatment as in the control population (**Fig. S6B**). While the trans-factors included in our guide panel did not appear to systematically regulate tRNA abundance, we reasoned that tRNA-internal cis regulatory elements (ICRs) could explain this stress-responsive effect. However, prior work assessing Pol III occupancy of tRNA genes across the human genome had not revealed obvious sequence correlations (*50*), though later reports found relevant differences of A and B box sequences to underlie differential tRNA gene occupation (*51*). While the global A and B boxes across all tRNAs are highly conserved at several positions (**Fig. 6G, S6F**), we observed that the 42 treatment-related upregulated tRNAs (**Fig. 6F**) yielded a well-defined motif within the center of the A box but a less distinct motif in the B box (**Fig. 6H, S6G, Methods**). This suggests that intrinsic sequence variation in the tRNA A box regulates stress-responsive transcription, a response conserved across mTOR and AAd perturbations.

Further relating A box motif enrichment to the guide-induced changes in the untreated ISR guide-transduced cells, revealed a positive correlation between the fold changes and the A box motif enrichment scores of individual tRNAs (ρ = 0.43, *p* = 0.004, **Fig. 6I**), supporting that the internal promoter motifs may partially explain the preferential transcription through the Pol III machinery under stress conditions. Transcriptionally, the stress cluster exhibited an upregulation of key integrated stress response genes (**Fig. S6I**) and concomitant changes in associated pathways (**Fig. S6J**), consistent with genetically induced stress phenocopying the tRNA pool changes observed with treatment. To further test whether tRNA genes differ in their dependence on trans-acting factors, we correlated the A box motif score of each tRNA gene with its expression fold change across perturbations (**Fig. S6K**, **Methods**). *TFIIIC* knockdown produced positive correlations in untreated cells (*GTF3C1*: ρ = 0.44, *p* = 8.4×10^-3^; *GTF3C6*: ρ = 0.34, *p* = 6.5×10^-3^), indicating that stronger A box genes are relatively spared, fitting the known stability of the TFIIIB-DNA complex, which sustains reinitiation once assembled (*52*). *POLR3C* knockdown had limited effect on A box motif scores in untreated or AAd cells, but showed a strong negative correlation under Torin-1 treatment (ρ = -0.84, *p* = 1.5×10^-4^). *POLR3C* encodes RPC3, which mediates the Pol III-TFIIIB contact disrupted by MAF1 upon mTOR inhibition, and as MAF1 is preferentially recruited to highly transcribed loci, we reason that the effect may scale with a context-dependent promoter strength.

Our findings corroborate previous studies reporting a functional asymmetry of the internal promoter sequences of A and B boxes (*53*), whereby conserved B box anchoring permits scanning of the Pol III complex across low-affinity A boxes (*54*). While the composition of the tRNA pool may underlie higher-order regulation under resting conditions, limited transcriptional resources under stress promote selective reprogramming of the tRNAome toward an optimized A box motif (**Fig. 6J**). We reason that the A box region may harbor interaction sites for poorly characterized factors regulating tRNA transcription, that a more comprehensive perturbation panel could reveal.

## Discussion

A flagship goal of 21st-century biology is creating a virtual cell that can simulate the function and behavior of cells in complex settings (*55*). Currently, substantial resources are being allocated to modeling high-dimensional data that couples scalable perturbations to cellular transcriptomes for sampling the degree of heterogeneity within a human cell. While scaling laws in other domains have suggested that ample data coupled with emerging generative artificial intelligence models may produce models with emergent properties, the complexity of this problem warrants multi-modal measurements of cells to predict the impact and fate of molecules (*56*, *57*). Because discordance between RNA and protein abundance has been widely noted in different contexts (*41–43*), we reason that a foundation model that accurately predicts mRNA abundances may often lack the context needed to determine a cell’s functional state, which is enacted primarily through proteins. Here, we introduce a new assay, sctRNA-seq, that is sensitive and specific to changes in tRNA expression from perturbation, differentiation, and stimulation. Our analyses show that for proteins like transcription factors and selenoproteins, concomitant quantification of tRNA species can resolve discordance between mRNA and protein levels, reinforcing the cellular tRNA pool as a dynamically regulated mechanism for protein translation.

While other technologies have nominally captured tRNAs at single-cell resolution (*18*, *58*, *59*), the limited information content hindered analogous insights revealed by our analyses. Though a probe-based detection of tRNA limits the one-to-one mapping of sequenced molecules to genomically-encoded tRNAs, our combined approach of probe annealing and gapfilling retains a feature-rich annotation of the cellular tRNA pool with sensitivity and depth to unveil previously undescribed principles of tRNA regulation. Additional measures of tRNA charging and post-transcriptional modifications (*60–62*) may further reveal the regulatory logic underlying cellular phenotypes that emerge from proteins encoded in mRNA and translated by tRNAs.

Mapping mRNA and tRNA changes to perturbation is a capacity uniquely afforded by our modular assay. However, the Flex assay retains broad utility for profiling tRNA abundances in arbitrary complex cell mixtures and tissues. These capabilities may further refine our emerging understanding of a protein regulatory logic encoded by tRNAs. These include specific isodecoders modulating malignant processes in cancer and infection (*10*, *63*). Further, our analyses of continuous differentiation trajectories in an *in vitro* model of human hematopoiesis suggest that the unique regulatory logic of tRNAs is transcriptionally coupled to lineage-defining protein-coding genes. These results support a model where cell state-specific proteins can be coupled during well-defined differentiation trajectories to establish functional proteomes for cell-type-specific function. While this transcriptional coordination was a feature of hematopoietic differentiation, our additional analyses show that complex regulatory principles similarly govern individual tRNA species, including the underlying sequence composition, which is variably responsive to transcriptional regulation by Pol III-mediated factors that are, in turn, induced by cellular stressors, including amino acid deprivation and mTOR perturbations. As a high-throughput perturbation-screening method for transcriptional and translational dynamics, we expect sctRNA-seq in additional studies, such as genome-wide screens, to reveal these tRNA transcription factors. Together, our work charts the widespread tRNA changes in heterogeneous conditions that reveal a previously underappreciated layer of cellular regulation.

## Supplementary Figures

**Figure S1.**
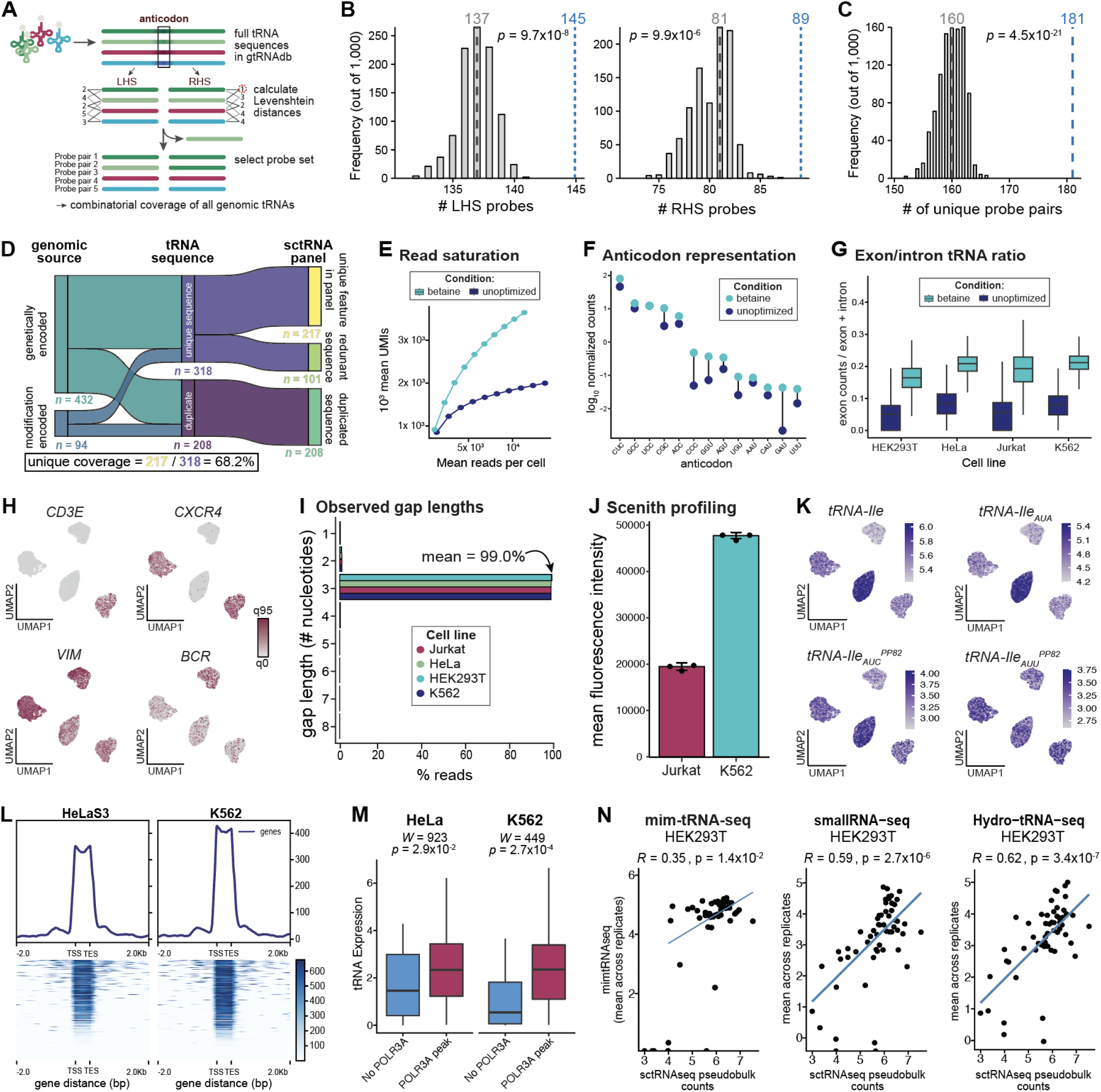
Supporting information for assay design and development. **(A)** Schematic displaying the probe panel design process, whereby the 25 bases left and right of the anticodon of all high-confidence tRNA sequences in the gtRNAdb are used to construct a reference. Probe candidates are selected to maximize the number of available sequences while ensuring a Levenshtein distance of at least 2 between every pair of probes. **(B)** Number of LHS (left) or RHS (right) probes after panel reduction through Levenshtein-distance filtering. The blue number marks the count of the rationally chosen probe set, applying an algorithm for the highest possible probe count. **(C)** Number of combinatorial probe pairs resulting from panel selection. The blue number marks the count of the rationally chosen probe set, using an algorithm to maximize probe count. **(D)** Sankey diagram tracing tRNA genes from genomic source (genetically vs. modification-encoded) through transcript sequence classification (unique vs. duplicate) to their representation in the sctRNA-seq panel (unique-in-panel, redundant, or duplicated). **(E)** Read saturation curves showing UMIs per cell as a function of downsampled read depth for the standard and betaine-containing gapfill conditions. **(F)** Per-anticodon normalized counts (log_10_) for the standard vs. betaine-containing gapfill conditions across non-encoded anticodon tRNA species, ordered by abundance. **(G)** Boxplots of exon/(exon + intron) tRNA ratio across all four cell lines (HEK293T, HeLa, Jurkat, K562) for both conditions. **(H)** UMAP embeddings of the four cell line mixture colored by marker gene expression (CD3E, CXCR4, VIM, BCR). **(I)** Distribution of observed gap lengths (in nucleotides) as a function of cumulative % reads, per cell line. **(J)** Mean fluorescence intensity from SCENITH profiling in K562 and Jurkat cells. Individual dots are replicates; bars are +/- error. **(K)** UMAP embeddings colored by amino acid-, isoacceptor-, or isodecoder-level tRNA-Ile expression. **(L)** Composite profiles and heatmaps of POLR3A ChIP-seq signal across tRNA genes in HeLa-S3 and K562, shown over a ±2000bp window around the tRNA gene body. **(M)** Pseudobulk expression of tRNA genes, split by POLR3A occupancy (red, occupied; blue, non-occupied). Occupancy was defined from POLR3A ChIP-seq in the matched cell line (HeLa-S3 for HeLa, K562 for K562). n = 33 vs. 76 genes (HeLa; non-occupied vs. occupied) and 21 vs. 88 (K562); P = 0.029 (HeLa) and P = 0.0003 (K562), two-sided Wilcoxon rank-sum test. Boxes span the interquartile range with the median indicated; whiskers extend to the most extreme values within 1.5x the interquartile range of the box. **(N)** Pairwise comparisons of per-anticodon expression estimates from mim-tRNA-seq, small RNA-seq and hydro-tRNA-seq against pseudobulk per-anticodon sctRNA-seq in HEK293T. Bulk method replicates are averaged using the mean of log_10_-transformed expression (n = 2 for mim-tRNA-seq; n = 3 for small RNA-seq and hydro-tRNA-seq); sctRNA-seq pseudobulk expression values are log_10_-transformed. Blue lines show linear fits. Spearman’s ρ and the corresponding p-value are given for each comparison.

**Figure S2.**
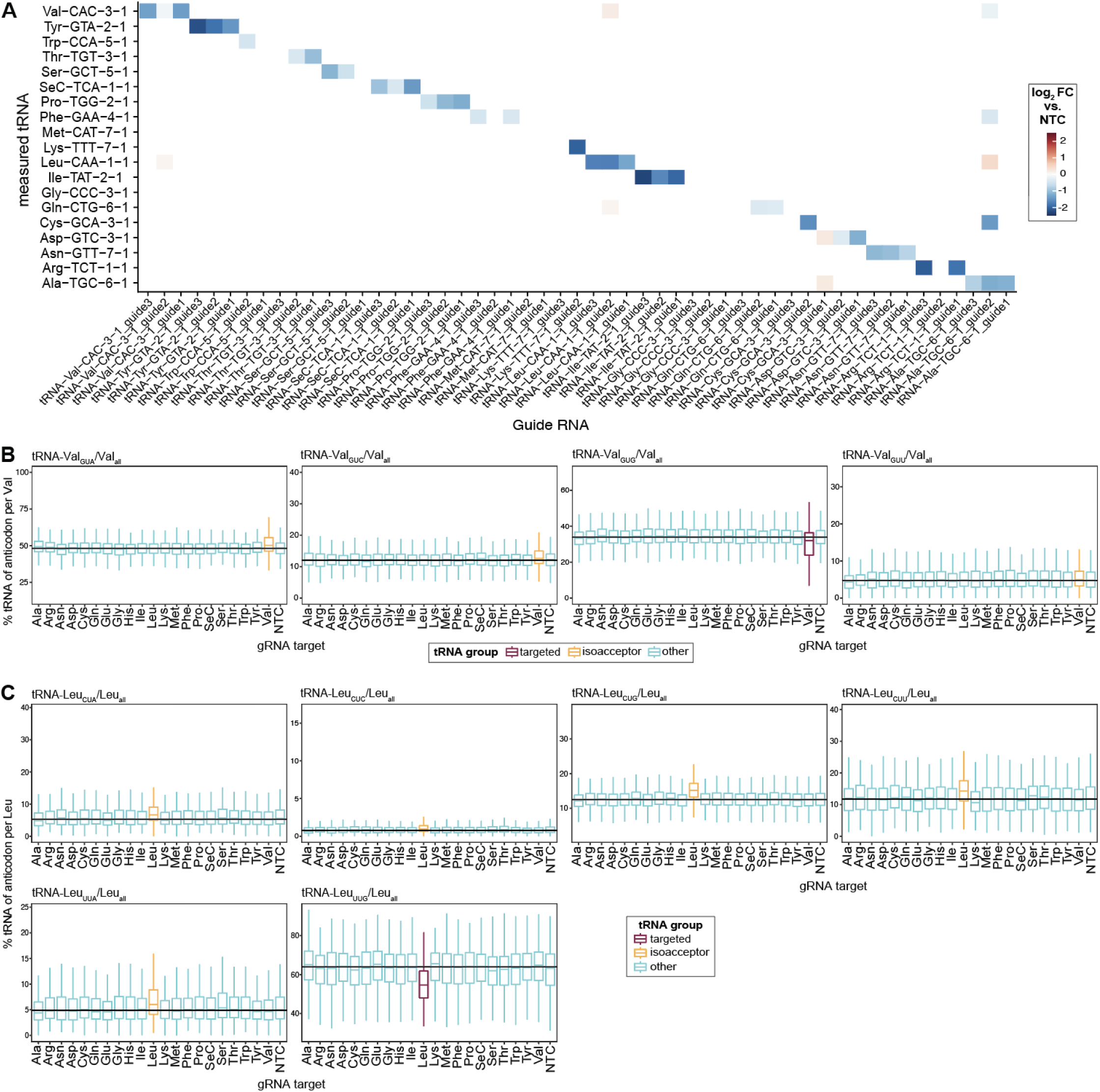
Supporting information for tRNA-targeting perturbations. **(A)** Heatmap showing the log_2_ fold change in tRNA abundance of individual isodecoders in response to single guides against NTCs. **(B)** Fractional abundance of each anticodon within the Val tRNA isoacceptor family (% isodecoder per isoacceptor), colored by whether the respective anticodon is the guide’s on-target isodecoder (red), on-target isoacceptor but off-target isodecoder (yellow), or not (false). **(C)** Fractional abundance of each anticodon within the Leu tRNA isoacceptor family (% isodecoder per isoacceptor), colored by whether the respective anticodon is the guide’s on-target isodecoder (red), on-target isoacceptor but off-target isodecoder (yellow), or another species (blue).

**Figure S3.**
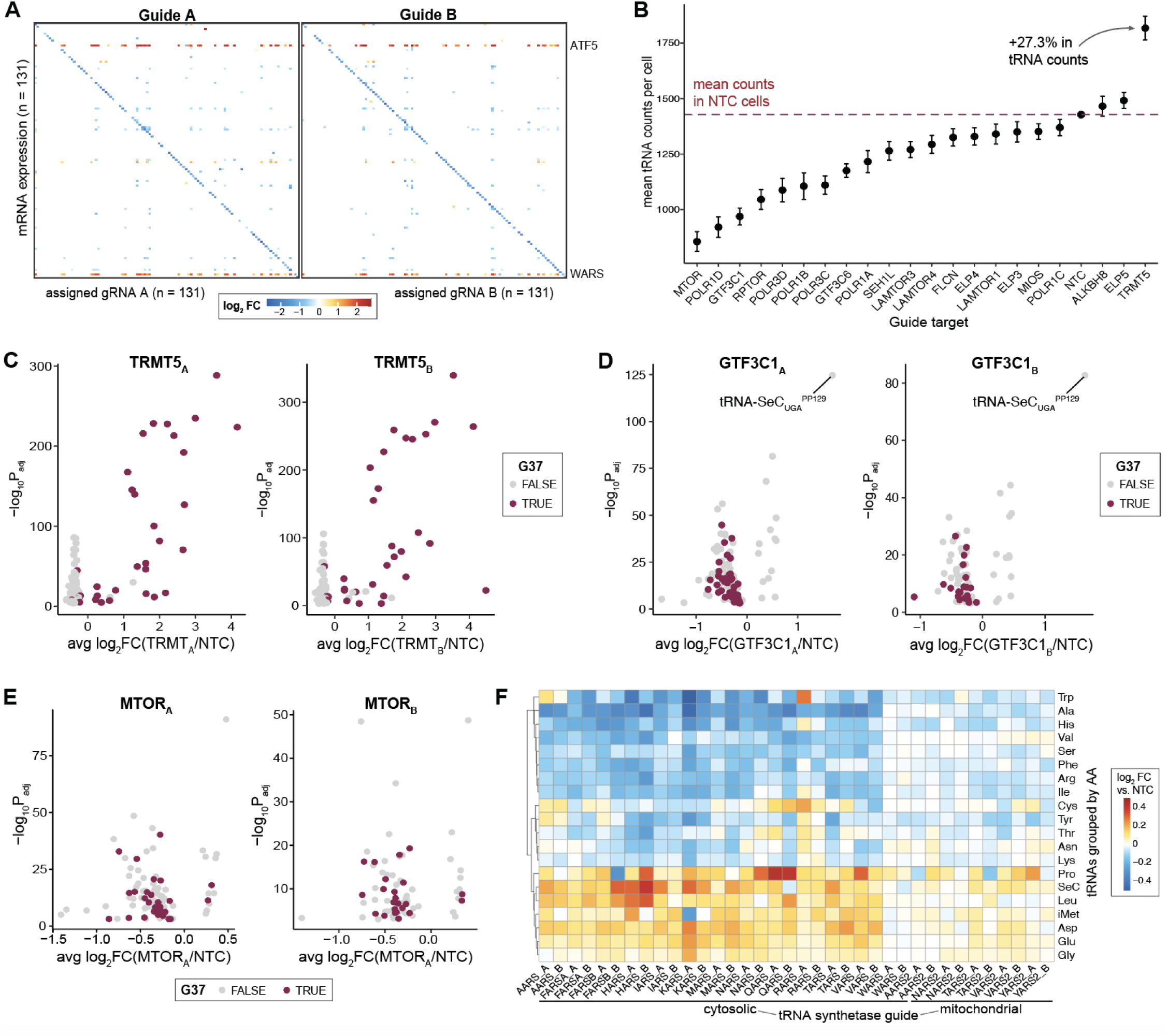
Supporting information for tRNA variability from mRNA-targeting perturbations. **(A)** Heatmap of mRNA expression across guide perturbations. The diagonal represents corresponding pairs of perturbation target and target gene expression. **(B)** Mean total tRNA counts per cell across tRNA-modifying, Pol III and mTOR-related guide targets. **(C)** Volcano plots of tRNA changes induced by guides A and B targeting TRMT5, colored by the identity of the nucleotide in position 37 of the tRNA molecule as either guanine (substrate of TRMT5) or non-guanine (not TRMT5 substrate). **(D)** Volcano plots of tRNA changes induced by guides A and B targeting GTF3C1, colored by the identity of the nucleotide in position 37 of the tRNA. **(E)** Volcano plots of tRNA changes induced by guides A and B targeting MTOR across guides targeting chromatin factors, colored by the identity of the nucleotide in position 37 of the tRNA. **(F)** Heatmap of log_2_ fold changes for tRNAs grouped by amino acid identity across tRNA synthetase-targeting guides.

**Figure S4.**
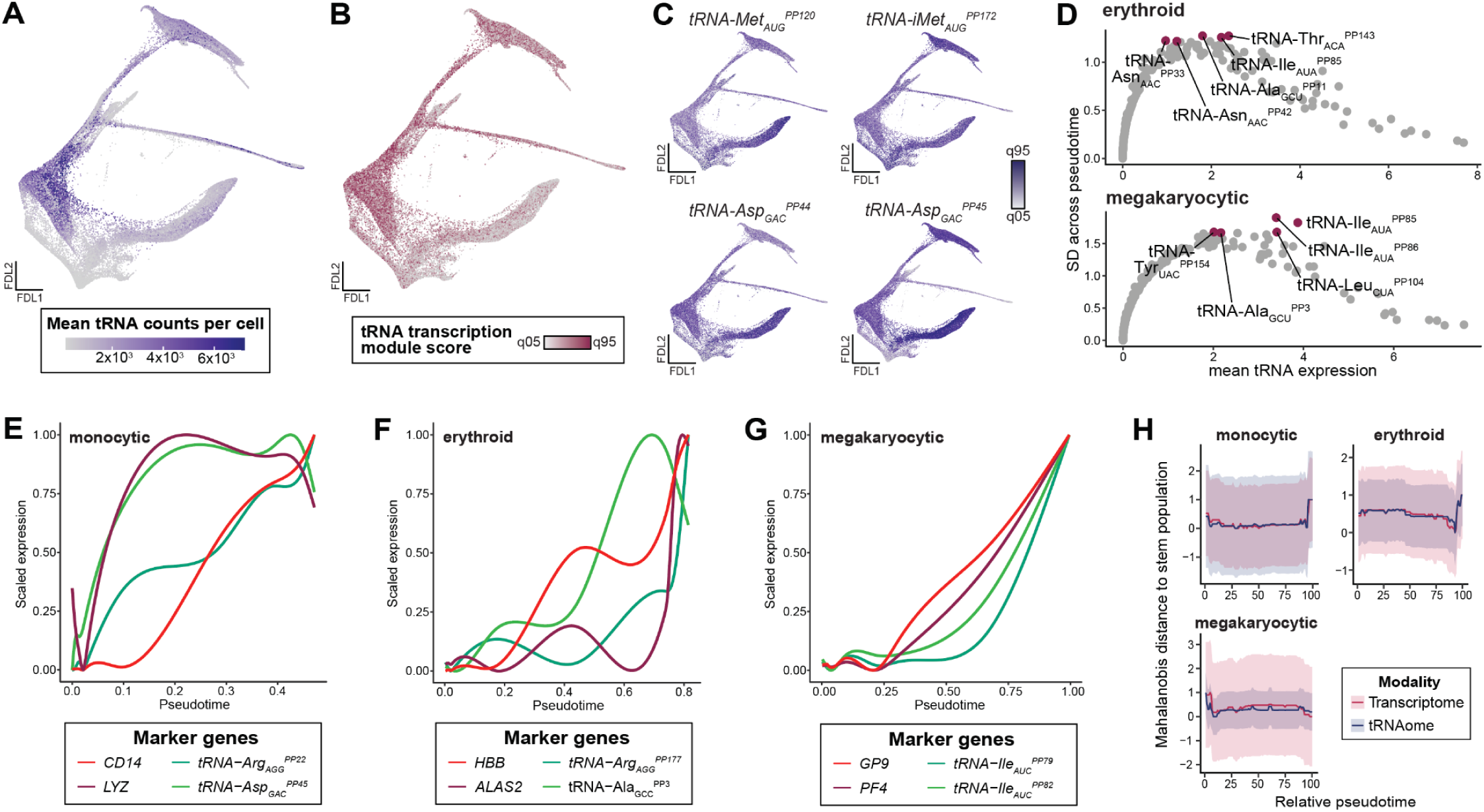
Supporting information for *in vitro* hematopoietic differentiation. **(A)** Force-directed layout colored by tRNA counts per cell. **(B)** Force-directed layout colored by UCell module score for the expression of components belonging to the tRNA transcription machinery. **(C)** UMAP colored by the expression of individual tRNA markers. **(D)** Scatter plot of expression and variability as mean standard deviation of individual tRNA markers in differentiating erythroid (top) and megakaryocytic (bottom) cells, highlighting the top 5 most variable tRNAs. **(E)** Trajectories of mRNA and tRNA markers over pseudotime for the monocytic cell population. **(F)** Trajectories of mRNA and tRNA markers over pseudotime for the erythroid cell population. **(G)** Trajectories of mRNA and tRNA markers over pseudotime for the megakaryocytic cell population. **(H)** Mahalanobis distance trajectories for each lineage computed with data permutation.

**Figure S5.**
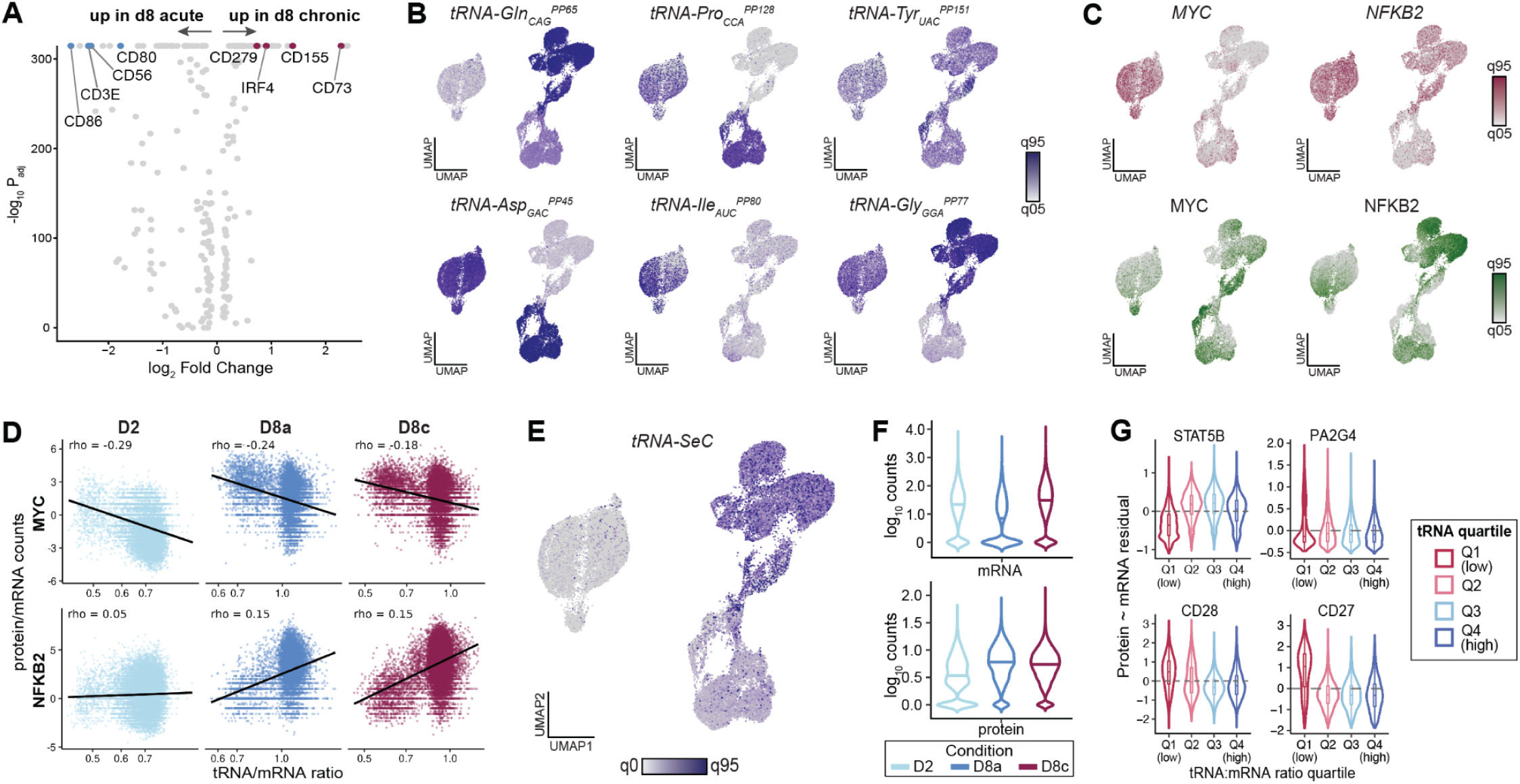
Supporting information for *in vitro* T cell stimulation. **(A)** Differential protein abundance between day 8 acute (D8a) and day 8 chronic (D8c) stimulation. Selected proteins are labeled; blue, enriched in D8a; red, enriched in D8c. **(B)** UMAP embeddings colored by expression of selected tRNAs. Subscripts denote the decoded codon; superscripts denote [feature identifiers]. Color scales span the 5th to 95th percentile. **(C)** UMAP embeddings coloured by *MYC* and *NFKB2* mRNA (top, red) and protein (bottom, green) expression. **(D)** Correlation of tRNA/mRNA ratio (x-axis) and ratio between protein and mRNA counts for MYC (top) and NFKB2 (bottom), split by condition. **(E)** UMAP embedding coloured by tRNA-SeC expression (0th to 95th percentile). **(F)** Distributions of *GPX4* mRNA (top) and protein (bottom) counts in D2, D8a and D8c. **(G)** Residuals of protein abundance after regression on mRNA, for STAT5B, PA2G4, CD28 and CD27. Cells are stratified into quartiles of total tRNA:mRNA ratio (Q1, lowest; Q4, highest).

**Figure S6.**
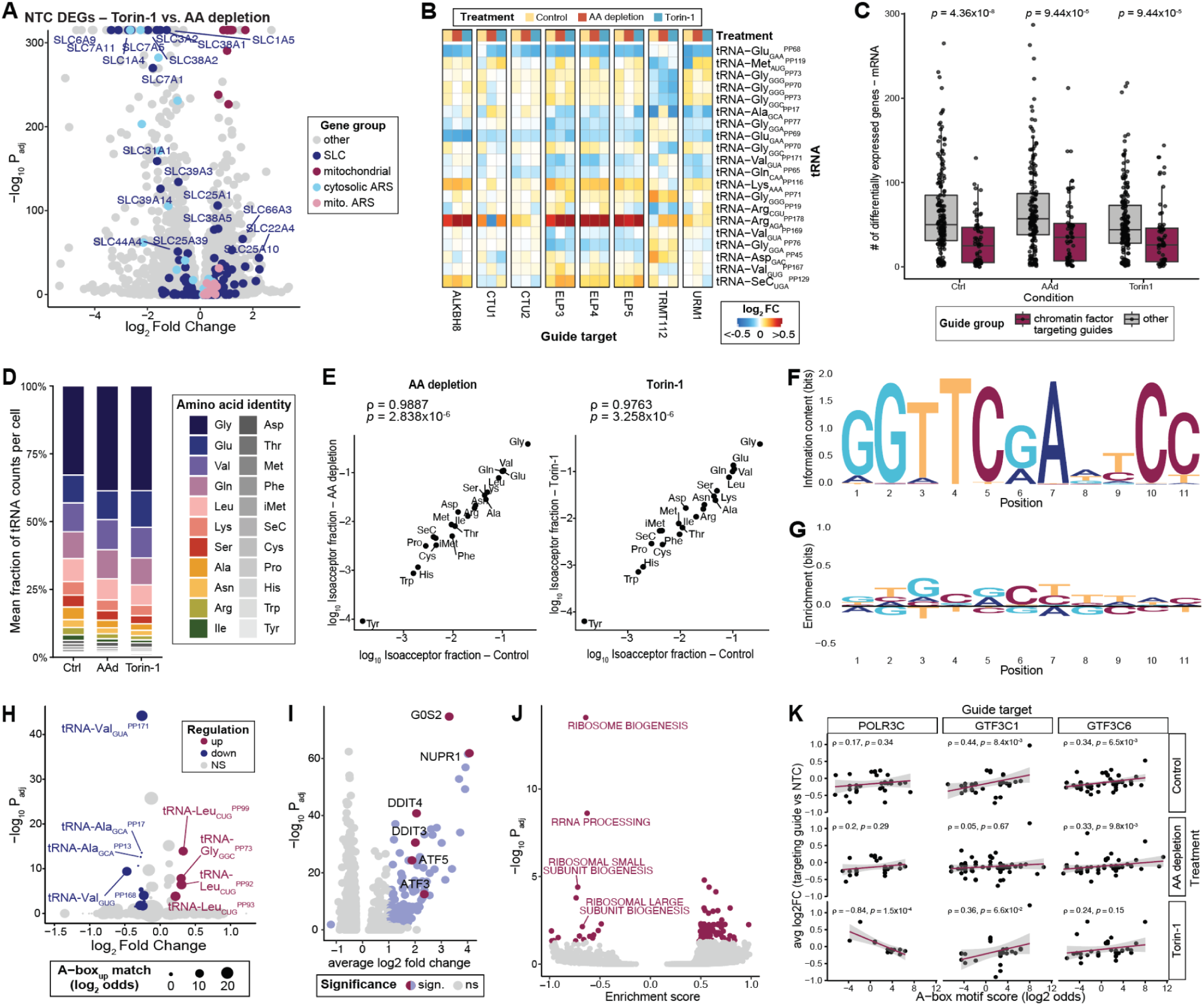
Transcriptional divergence and tRNAome convergence between amino acid depletion and Torin-1 treatment. **(A)** Differentially expressed genes between Torin-1 and amino acid depletion in NTC cells. Positive log₂ fold change indicates relatively higher expression in AA depletion. Highlighted groups: solute carrier (SLC) transporters, mitochondrial genes, cytosolic aminoacyl-tRNA synthetases (ARS), and mitochondrial ARS. Selected SLC genes are labeled. **(B)** Heatmap of tRNA abundance changes in cells carrying guides targeting tRNA-modifying enzymes (columns), relative to NTC within each treatment condition. **(C)** Number of differentially expressed mRNAs per perturbation, grouped by whether the guide targets a chromatin remodeling factor (red) or any other guide class (grey), across the three treatment conditions. Each dot corresponds to one guide target. **(D)** Mean fraction of tRNA counts per cell in NTC cells by amino acid identity for each treatment condition (compositional version of Fig. 6D). **(E)** Isoacceptor tRNA fractions under amino acid depletion (left) or Torin-1 treatment (right) compared to the untreated control. **(F)** Top: Sequence logo of the B box motif across the whole tRNAome. Letter height indicates information content (bits). Bottom: Enriched sequence motifs of the B box internal promoter elements of the 42 tRNA isotranscripts significantly upregulated in Fig. 6F. Letter height indicates enrichment (bits). **(G) (H)** Volcano plot of differential tRNA isotranscript abundance for untreated cells transduced with guides belonging to the group clustering with the treatment-condition guides in (C) tested against untreated NTC cells. Significant isodecoders (adjusted p-value < 0.05, log_2_ FC > 0.1) are shown in red (upregulated) and blue (downregulated). Point size indicates log₂ odds of an A-box match to the upregulated-tRNA motif in (Fig. 6H). **(i)** Differentially expressed genes for untreated cell population, testing guides that cluster with treatment conditions in Fig. 6C against untreated NTC guides. Significant genes are colored. **(J)** Gene set enrichment analysis of untreated cell population, testing guides that cluster with treatment conditions in Fig. 6C against untreated NTC guides in GO Biological Process. Negatively enriched terms feature predominantly ribosome processing-associated pathways, such as ribosome biogenesis and rRNA processing. **(K)** Correlation plots for A box motif score and log_2_ fold change of differentially expressed tRNAs for *POLR3C*-, *GTF3C1*-, or *GTF3C6*-perturbed cells across treatments against NTC guides for the respective treatment.

## Materials and Methods

Our research complies with all relevant ethical and regulatory guidance, including the Institutional Review Board at Memorial Sloan Kettering Cancer Center. Our project was exempt from human subjects research.

### Design of sctRNA-seq probes

Due to the high homology among tRNAs, we developed a custom heuristic for tRNA probe designs to establish our tRNA panel. Explicitly, we define the set of genomically encoded tRNA loci as a set *T*. Our goal is to assemble two sets of 25 nucleotide probes, *L* and *R*, which would hybridize the 25 nucleotides on the left-hand side (LHS) and the 25 nucleotides on the right-hand side (RHS) of the anticodon for all individual tRNAs, *t*. For simplicity, we denote *t_LHS_* and *t_RHS_* as the reverse complement of the 25 bases on the LHS and RHS, respectively (noting the reverse complement would be the ssDNA probe used for annealing). We use a Levenshtein distance function *d* to account for mismatch tolerance in the ssDNA probes. To identify the probes for *L* and *R*, we imposed the following constraints ∀ *t* ɛ *T*, ∃ *l* ɛ *L* ∧*r* ɛ *R* s.t.:

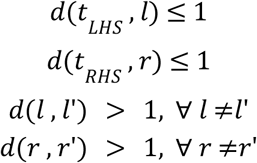

These constraints require each probe to be at most a Levenshtein distance of 0 or 1 from the reverse complement of each tRNA locus. Further, each LHS or RHS probe must be at least a Levenshtein distance of 2 from every other probe in its respective set. While ssDNA probes of length 25 can likely tolerate mismatches beyond 1 nucleotide depending on the position and nature of the change, we reason that enforcing at least two differences between all probes in the panel would, on average, hybridize the closest sequence to the underlying tRNA even in the presence of post-transcriptional modification outside of the backbone.

We used a greedy algorithm (i.e., local best choice) to define *L* and *R* to maximize the number of individual probes in both sets, thereby optimizing the number of tRNAs that could be uniquely identified under constraints of sequence specificity while tolerating one mismatch (due to sequencing error or variable hybridization efficiencies. Specifically, for a possible *t_LHS_*, we selected a probe (*l*) to be appended to the set (*L*) such that the choice of *l* had the lowest similarity to the fewest unaccounted-for tRNAs (| *d*(*t_LHS_*, *l*) ≤ 1 |). We repeated the same procedure for *r*, *R*, and *t_RHS_*. R code to reproduce this heuristic is provided as part of our GitHub resource noted in the **Code Availability**. To complement this algorithmic determination of tRNAs, we further appended rationally designed probes targeting the 32 species of intron-containing tRNAs to include 3 total pairs per tRNA: one constant RHS and two variable LHS such that each one would hybridize variably whether the mature (spliced or intron-removed) species was present or the immature form (intron-containing).

### Gapfilling method and development

We designed our sctRNA-seq assay to enable multi-modal profiling of transcriptional and translational dynamics in single cells based on the Chromium GEM-X Flex workflow (10x Genomics), using paired custom probes combined with an enzymatic gapfill (**Fig. 1a**) reaction for tRNA isotype identification. The principles of the rationally designed ssDNA probes paired with a short gap-filling reaction were previously described as the Genotyping In Fixed Transcriptomes (GIFT) method (*21*). Here, primary method development focused on optimal design of a tRNAome-targeting probe set and optimization of experimental conditions. For the experimental validation, the primary heuristics for assay development was a characterization of the resolution of tRNA molecules (e.g., at the species or isodecoder level), maturation stage (e.g., pre- or post-transcriptional modifications; with or without retained introns), and the enzymatic gapfilling efficiency. Similar to GIFT, we verified that the addition of betaine in the gapfilling reaction yielded substantially more tRNA counts, which is consistent with its previously reported ability to mitigate GC-dependent secondary structure formation across duplexes (*64*). Indeed, we observed an increase in recoverable library molecule counts and included betaine in all subsequent experiments.

### Cell fixation, permeabilization, and storage

We fixed and permeabilized cell lines and primary cells as described in the manufacturer’s sample preparation handbook (CG000782). Briefly, cells were washed in 0.05% PBS with BSA (Miltenyi Biotec, 130-091-376), centrifuged at 300 rcf for 5 min, upon which supernatants were removed and the cell pellets were resuspended in Fix & Perm buffer as prepared according to the manufacturer’s instructions. Cell suspensions were incubated at 4°C for 24h, after which Additive C (in case of 10x Genomics Flex v2) was added, and centrifuged at 850 rcf for 5 min. Supernatant was removed and cell pellets were resuspended in 1 mL Quenching buffer. For improved cell permeability, samples were subjected to one freeze thaw cycle according to the long term storage instructions in the sample preparation handbook. Briefly, 100µL of Enhancer (10x Genomics, 2000482) and 275 µL of 50% glycerol were added to 1 mL of cell suspension and stored at -80°C upon further usage. For profiling, frozen cell stocks were thawed at RT, centrifuged at 850 rcf for 5 min and resuspended in 1 ml of Quenching buffer. Cells were stored on ice until further usage.

### 10x Genomics Flex

The microfluidics-based single-cell workflow and library preparation were performed according to the 10x Genomics Flex protocol, following the manufacturer’s instructions with relevant modifications as published previously to extend the protocol by the gapfilling reaction (*21*) and as outlined in **sctRNA-seq method and development**. Briefly, depending on the sample species, we hybridized either human or mouse 10x Genomics whole transcriptome probes with human or mouse tRNAome probes at 32 nM. For perturbation experiments, we added an additional guide-capture probe panel at 32 nM. After hybridization and post-hybrization washes, cells were counted and 120K cells were aliquoted and resuspended in Gapfill enzyme mix with Betaine (except for a comparative cell line mixing experiment as pointed out above). The cell mixture was incubated at 25°C for 10 min and 37°C for 20 min, after which we performed one wash step. Subsequently, cells were counted again, and the required cell count was directly used as input for the Chromium chip. Raw basecall files (.bcl/.bases) were converted to .fastqs using bcl2fastq (Illumina) or bases2fastq (Element) for further processing using CellRanger v8 (10x Genomics). We quantified WTA libraries against the human probe set v1.1.0 using default parameters.

### Guide RNA design, selection, construction, and screening

Guide RNAs targeting tRNA genes were designed using the CRISPick tool (*65*). Per target gene, the three highest-scoring guide designs were selected by requiring a GC content between 0.3 and 0.7 that were at least 10 bases apart from the ranked gRNA designs. For protein-coding gene perturbations, guide RNAs were selected from previously designed gRNAs with demonstrated transcriptional activity (*31*). Guide targets were rationally selected to enrich for pathways associated with tRNA biosynthesis or post-transcriptional modifications based on previous supervised clustering. Additional chromatin-modifying guides were appended for specific hypothesis testing of the role of chromatin modifying enzymes in tRNA transcriptional regulation. NTC guides were selected based on minimal transcriptomic perturbation effects in previous screens.

All sgRNA libraries were ordered as oPools from Twist with 5’ overhang 5’-agtatcccttggagaaccaccttgttg-3’ and 3’ overhang 5’-gtttaagagctaagctggaaacagcatagcaag-3’ and cloned into the backbone via Gibson assembly as previously described (*66*). Briefly, the backbone pCRISPRia_v2 (Addgene #84832) was enzymatically digested in two consecutive steps using BstXI (NEB, R0113L), followed by gel purification from a 1% agarose gel, and BlpI (NEB, R0585L). After DNA purification using the NEB monarch DNA purification kit according to the manufacturer’s instructions (NEB, T1030S). sgRNA oPools were resuspended in IDTE buffer and amplified via PCR with 2x KAPA HiFi Master Mix (Roche, 07958935001) using primers P276F and P276R. The reaction was run in a thermal cycler with the following parameters: 95°C for 3 min, 64°C for 15 min, 72°C for 10 min, 4°C hold. After purification of the double-stranded product (NEB, T1030), the library was cloned into digested pCRISPRia_v2 via HiFi assembly (NEB, E2621) for 1 hr at 50°C, using a 5:1 molar ratio of insert library to vector. The resulting vector library was electroporated into Endura Electrocompetent Cells (LGC, 60242-2) with an Eppendorf Eporator per manufacturer protocol. After electroporation, each reaction was incubated in 1 ml LGC Recovery Media at 32°C for 1.5 hr, then expanded into 100 mL LB + ampicillin overnight culture. 30mL of culture was prepared via miniprep (Qiagen, 27104).

Lentivirus was produced by transfecting HEK293T cells (ATCC, CRL-3216) with the lentiviral helper plasmids VSVG (Addgene #8454) and psPAX2 (Addgene # 12260), sgRNA library, and polyethyleneimine (PEI) (Kyfora Bio,23966100). For one 15 cm dish of HEK293T 20 µg of plasmid DNA, 10 µg of VSVG, 15 µg of psPAX2, and 180 µL of 1 mg/mL PEI were mixed in 2ml of Opti-MEM and added to cells. Media was changed 12 hours post-transfection. The media containing virus was collected 48 hours post-transfection and filtered using 0.45 µm SFCA filters (Corning).

K562 ZIM3-dCas9 cells (*67*) were a gift from Tom Norman. Lentivirus was titrated to determine optimal concentration for an MOI of 0.1, based on BFP signal. For lentiviral transduction, K562 ZIM3-dCas9 cells were transduced at 200x coverage in 6 well plates. 1 million cells were combined with an appropriate amount of virus and media was added to 2 mL. 4 µg/mL polybrene (Sigma, TR1003G) was added to cells which were spininfected at 800g for 45 mins. Media was changed 24 hours post-transduction. 48 hours post-transduction cells were sorted on BFP. On day 7, cells were fixed as described above and stored at -80°C.

### Cell lines and cell culture

In our first experiment to study tRNA expression variability, we established our protocol on a mixture of human cell lines. Specifically, HEK293T, HeLa, Jurkat, and K562 lines were procured from ATCC and monitored for mycoplasma contamination. Genotyping at ATCC confirmed the identity of all lines. Analyses of expression in **Figure 1** matched known RNA-seq profiles for these lines, including established marker genes *CD3E, CXCR4, VIM,* and *BCR*. For SCENITH profiling, 1,000,000 cells in 1 mL of cell culture media were plated in one well of a 6 well culture dish. BFA (Biolegend, 420601) or BFA and Puromycin (BFA+Puro, Gibco, A1113803) were diluted 1:100 in complete culture media. 100 µl of either BFA or BFA+Puro were added to each well. Cells were then incubated for 20 mins in the incubator. After 20 minutes, cells were collected and washed twice with PBS+BSA (10% PBS with 0.5% BSA, Fisher Scientific BP1600-100). Cells were then stained with a viability dye (Invitrogen, L34963) according to manufactures instructions, and washed twice with PBS+BSA. Cells were then fixed and permed (eBioscience, 00-5523-00) according to manufacturers instructions. Blocking buffer consisting of TruStain FcX Blocker (Biolegend, 422302) and True-Stain Monocyte Blocker (Biolegend, 426102) diluted 1:100 was made and 50ul was added to cells for 15 mins. Anti-Puromycin antibody (Sigma Aldrich, MABE343-AF647) was diluted 1:100 in 50 µl blocking buffer and cells were stained for 30 mins on ice. After 2x washes in permeabilization buffer cells were analyzed on the Attune flow cytometer.

### Primary cell cell culture

Primary CD34+ cells were purchased from STEMCELL Technologies. The cells were thawed and expanded for 5 days at a density of 150k cells in 15 ml StemSpan media supplemented with CC100. On day 0, the culture was split up into three equal parts for further differentiation. StemSpan Erythroid Expansion Supplement (STEMCELL, 02692), Megakaryocyte Expansion Supplement (STEMCELL, 02696), and Myeloid Expansion Supplement II (STEMCELL, 02694) were added, respectively, and supplemented every two days until day 7. Cells of each culture were sampled, washed in PBS with 0.05% BSA and up to 10M cells were fixed as described above.

### Human T-cell isolation and activation

Human peripheral blood mononuclear cells (PBMCs) were isolated from human buffy coats (New York Blood Center). Each 10-ml aliquot of buffy coat was first transferred to a 50ml Falcon tube, diluted with 25mL PBS, and underlaid with 10ml Accu-prep (Accurate Chemical). Samples were then centrifuged at 800g at 30°C for 20 minutes with acceleration and brake turned off. The white buffy coat layers containing PBMCs were collected, pooled into sterile Falcon tubes, washed with PBS, and centrifuged for 6min at 500g at 30°C with the brake on. Cell pellets were then resuspended in 5 mL ACK buffer (150mM Ammonium chloride, 10mM Potassium bicarbonate, 0.1mM Disodium EDTA) and incubated at room temperature to lyse red blood cells. After 10 minutes, the tubes were topped up to 50ml with PBS, centrifuged, and resuspended at 1-2 x10^8^ cells per ml in isolation buffer (PBS without calcium or magnesium, supplemented with 2% FBS and 2 mM EDTA). T-cells were then isolated with the Dynabeads™ Untouched™ Human T Cells Kit (Invitrogen 11344D) following manufacturer’s instructions and activated with plate-bound anti-CD3 (Biolegend 317349 (clone OKT-3), 5ug/ml) and anti-CD28 antibody (Biolegend 302944, 2ug/mL) for 2 days under RPMI-1640-based T-cell media (10% FBS, 2mM L-Glutamine, penicillin-streptomycin, 50µM 2-Mercaptoethanol, 5 ng/ml of human IL-7 (Peprotech 200-07), and 10 ng/ml of human IL-15 (Peprotech 200-15) at a density of 2×10^6^ cells per ml.

After 48 hours of T-cell activation described above, T-cells are replated in 6-well plates pre-coated with (chronic) or without (acute) anti-CD3 antibody (Biolegend 317349 (clone OKT-3), 5ug/ml) at a density of 1×10^6^ T-cell per ml in T-cell media as described above. For every 48 hours, cells are resuspended, spun down, and re-plated with fresh T-cell media with the respective stimulating conditions. Chronically stimulated or time-matched acutely stimulated T-cells are harvested for experiments at the time stated in each experiment.

### Protein labelling for trimodal T cell profiling

CITE-seq antibody panels were provided by Proteintech (Proteintech, G900150). The antibody preparations were reconstituted and mixed as instructed by the manufacturer (Proteintech, C10013). Briefly, antibody cocktails were equilibrated at RT for 5 min and spun down at 1.5kxg for 30 s. Reconstitution was performed by adding 52 µl BSA buffer (10% PBS + 1% BSA) to the Cell Surface Cocktail, while adding 55 µl of IC Staining Buffer (10% PBS, 1x RNase Inhibitor, 7.55% Nuclease-free BSA, 0.2% Tween-20, 0.5% NP-40, 0.025% Dextran Sulfate 8 kDa, 1x Enhanced Blocking Reagent (Proteintech, G900005), supplemented with 4.5 µl FcX) to the Intracellular Cocktail and Intracellular Supplement, respectively. After 5 min of incubation at RT, vortexed the antibody cocktails for 15 s and centrifuged at 10kxg for 30 s. The reconstituted cocktail was transferred into a 1.5 mL Eppendorf LoBind Protein tube (Eppendorf, 022431081) and centrifuged at 14kxg for 10 min, upon which 50 µl of the Cell Surface Cocktail was transferred into a fresh tube to remove potential protein aggregates. 53 µl of the Intracellular Cocktail and Intracellular Supplement respectively were combined into a new tube (“Intracellular Antibody Mix” hereafter). Antibody cocktails were stored on ice until further use.

Intra- and extracellular cell staining was performed for the extracellular and intracellular panel according to the manufacturer’s protocol, following the 2-wash option (10X Genomics CG000529). Briefly, T cells were removed from culture and washed in 10% PBS + 1% BSA. Cells were counted, followed by transfer of 1M T cells into a new tube, centrifugation at 400 rcf for 5 min and removal of supernatant. For cell surface blocking, the cell pellet was resuspended in 50 µl of PBS + 1% BSA and 5µl of Human TruStain FcX (BioLegend, 422301) was added for incubation at 4°C for 10 min. 50 µl of the Cell Surface Cocktail was added to the Fc blocked cells and gently resuspended before incubation at 4°C without light exposure for 30 min. After incubation, 1.4 ml of 10% PBS + 1% BSA was added to the surface-stained cells and gently pipette-mixed. Samples were centrifuged at 400 rcf for 5 min at 4°C and supernatant was removed without disturbing the cell pellet. After resuspension in 1.5 ml of 10% PBS + 1% BSA, cells were spun down again (400 rcf, 5 min), resuspended in 1 ml of room temperature Fixation Buffer (1x Conc. Fix & Perm Buffer (10x Genomics, 2000517), 4% Formaldehyde), and incubated for 20min at RT. After centrifugation at 2000 rcf for 5 min at RT, cells were resuspended in 1 ml of Quenching Buffer. After centrifugation at 2000 rcf for 5 min at 4°C, supernatant was removed and the cell pellet was resuspended in 100 µl of the Intracellular Antibody Mix and incubated at 4°C for 30 min. After incubation, 1 ml of chilled Intracellular Wash Buffer (10% PBS, 0.2 U/µl RNase Inhibitor, 0.1% Tween-20, 1% BSA) was added and cells were centrifuged at 2000 rcf for 5 min at 4°C. Supernatant was removed and cells were submitted to sample fixation and storage until further profiling as described above.

### *In vitro* cellular stress model

On day 7 after transduction as described above, dCas9^+^ K562s were split into three cultures of 1M cells, spun down and resuspended in complete media (for the control and Torin-1 conditions) or amino acid dropout media (MSKCC media core). The mTORC1 inhibition sample was treated with 250 nM Torin-1 in DMSO, and equal volumes of DMSO were added to the control and AA depletion cultures. After 12h, cells were spun down and fixed as described above.

### Downstream analyses

#### Raw data pre-processing

Raw Fastq files were processed using CellRanger v8 following standard recommendations. We analyzed standard WTA mRNA libraries using CellRanger ‘count’ with the Human Probe Set v1.1.0 reference for all experiments. The custom tRNA libraries are separately amplified and indexed, resulting in a separate library (.fastq.gz) that was processed with *giftwrap* (*21*). At a high level, giftwrap parses the custom library to determine the cell barcode, UMI, identity of the tRNA-targeting RHS and LHS probes as well as the combination (probe pairing) thereof, and the anticodon-identifying gapfill product with accompanying quality metrics. When running giftwrap via the command-line interface, we match cell barcodes against a library-specific set of barcodes provided by the filtered feature-barcode matrix from CellRanger outputs for 10x Genomics Flex. We resolved PCR duplication by grouping reads with matching cell barcodes and UMIs within one edit distance. In rare instances where the same UMI matches multiple probe pairs in the same cell, we retain the most abundant to reduce potential artifacts from PCR recombination. Finally, a consensus call of gapfilled sequences is determined across all PCR duplicates for each cell/UMI pair. After filtering, the output files containing barcodes (barcodes.tsv), probe counts (probe_reads.tsv), and probe sequence mapping (manifest.tsv) are used for downstream analyses.

For gRNA and CITE-seq libraries, we parsed multiplexed .fastq files using cutadapt (*68*) before quantification with kallisto|bustools (*69*),(*70*) using the protospacer gRNA or antibody barcode dictionary. We constructed all indices using a 15-mer reference with up to 1 mismatch per barcode. We performed error correction and UMI collapsing following bustools best practices.

#### Construction of Seurat objects and integration of tRNA data

We read gene expression matrices from 10x-format HDF5 files. Cell barcodes were truncated to the 16-nt 10x barcode and appended with a sample identifier, and per-sample objects were merged into a single Seurat object (Seurat v*5.5.1*). tRNA probe-pair reads were quantified with *giftwrap* (v0.6.1), which returns per-read assignments of cell index, probe index and 3-nt gapfill sequence together with the corresponding barcode and probe manifests. We assigned each read to its cell barcode and probe pair, and annotated probe pairs with their intended amino acid using the human probe-pair design table (**Table S1**). Reads with incomplete annotation or with a gapfill sequence of length other than 3 nt were discarded. The filtered gapfill sequences were taken as the observed anticodon and converted to RNA nomenclature. Reads derived from different sample-identifying barcodes of the same library were combined at this stage.

Three count matrices were constructed by counting reads per cell and feature. The probe-pair matrix used the combination of probe pair, intended amino acid and observed anticodon as its feature, retaining only features supported by at least 25 reads across the dataset. The anticodon- and amino-acid-level matrices were built from the subset of reads passing an anticodon-validity filter: observed anticodons were matched to a curated anticodon-to-amino-acid reference (**Table S5**), reads whose anticodon was absent from this reference were discarded, and a read was retained only if the amino acid encoded by its observed anticodon was consistent with the amino acid of the probe pair. This criterion retains isoacceptor-consistent anticodons that are not themselves encoded in the genome, while removing probe-pair/anticodon combinations that are incompatible with the targeted isotype. In all three cases, only barcodes present in the mRNA object were retained. We added the three matrices to the Seurat object as separate assays (tRNA_pp, tRNA_ac, tRNA_aa), and removed cells with fewer than 50 total counts in any assay, so all retained cells carry paired mRNA and tRNA measurements at all three levels.

Finally, we filtered the probe-pair assay to valid features using the same anticodon criterion, and stored the filtered matrix as an additional assay (tRNA_pp_valid) used for all downstream analyses at the probe-pair level. As a quality control, we annotated observed anticodons by whether they are encoded in the human genome (gtRNAdb), and inspected total counts per anticodon by encoding status.

#### Data normalization, cell clustering and visualization

The whole transcriptome (“RNA”) and each tRNA assay was processed independently with the same workflow. Counts were log-normalized (*NormalizeData*, scale factor 10,000), variable features were selected by the variance-stabilizing transformation (*FindVariableFeatures*, up to 2,000 features; because all three assays contain fewer than 2,000 features, all features were retained), and normalized values were centered and scaled to unit variance (*ScaleData*). We performed principal component analysis on the scaled data (*RunPCA*). A shared-nearest-neighbor graph was constructed on the first *N* principal components (*FindNeighbors*) and cells were clustered using the Louvain algorithm at a resolution of 0.2 (*FindClusters*). For visualization, we computed UMAP embeddings from the first 30 principal components (*RunUMAP*) of the RNA assay in all figures, except when the embedding in tRNA space is indicated. Here, *we computed RunUMAP* with 30 principal components on the assay tRNA_pp_valid.

#### Cell line analyses

To assess sequencing saturation of the tRNA probe assay, per-read tables from giftwrap were annotated as described above: each read was assigned its cell barcode, probe pair and intended amino acid, and reads with incomplete annotation or a gap-fill sequence other than 3 nt were discarded. Reads were then restricted to barcodes called as cells in the corresponding gene expression library. We generated saturation curves by computational downsampling. For each target depth *d* ∈ {0.1, 0.2, …, 1.0}, we drew the number of reads retained for each unique molecule from a binomial distribution with size equal to that molecule’s observed PCR duplicate count and success probability *d*. We counted a molecule as detected if at least one of its reads was retained. The mean and median number of detected molecules (UMIs) per cell were recorded at each depth, together with the corresponding mean number of retained reads per cell, and mean UMIs per cell were plotted against mean reads per cell. Curves were computed independently for the betaine-free and betaine-containing conditions (4CL and 4CLB) for comparison on the same axes.

We inferred cell cycle phase for each cell using the built-in Seurat function *CellCycleScoring*, applied to log-normalized expression values. S-phase and G2/M scores were calculated from the canonical gene sets bundled with Seurat (cc.genes), derived from Tirosh et al.(*71*) to compute the mean expression of the gene set minus the mean expression of a control set of genes randomly sampled from expression-matched bins, following the *AddModuleScore* approach. We then assigned cells to S or G2/M based on the higher of the two scores, and to G1 when both scores were negative. Cell lines were identified by enrichment of defining signatures resolving HEK293T (i.e., *GATA1*, *LYZ*, *BCR*), HeLa (i.e., *NCAM1*, *TTC26*, *SPI1*), Jurkat (i.e., *CXCR4, CD3E, TRBC2*), and K562 (i.e., *S100A*, *CD44*, *KRT8*).

The distribution of gapfilled sequences was assessed by modifying the import function for the giftwrap-processed data above to retain features of all gapfill sequence lengths (instead of removing gapfill sequences of lengths other than 3 nt). This produced a feature x count matrix, which we added to the Seurat object as a separate assay. Counts were split by cell line, aggregated across cells for each feature, and grouped by gapfill length and summed up.

To compare sctRNA-seq, we obtained processed per-anticodon mim-tRNA-seq, small-RNA-seq and hydro-tRNA-seq read counts from Gene Expression Omnibus (GEO; GSE152621, GSE137834). Counts were medians of ratios-normalized and log_10_ transformed. Means were computed across technical replicates. Single-cell tRNA sequencing data generated through TotalX or Vasa-seq were obtained via GEO accessions GSE315939 and GSE176588. Processed POLR3A ChIP-seq tracks and peak calls for HeLa-S3 and K562 were downloaded from the ENCODE portal (Accessions ENCFF457PID, ENCFF392IDZ, ENCFF627DJP, ENCFF767DMU). Composite profiles and heatmaps were created using deepTools (3.5.6). We computed pseudobulk expression per tRNA and compared tRNA expression of those that overlapped a POLR3A peak to those that did not using a Wilcoxon rank sum test.

#### Differential expression analysis

Differential analyses of gene expression, tRNA features and other modalities where applicable were performed using the Seurat function *FindMarkers*. We grouped cells by metadata according to experimental context, and ran each comparison in a one-versus-rest design unless stated otherwise, testing the specified group against all remaining cells. We tested log-normalized values of the relevant assay using the default two-sided Wilcoxon rank-sum test, and adjusted p-values for multiple testing across all tested features using the Bonferroni method as implemented in Seurat. We considered features differentially abundant with an adjusted p-value below 0.05 and an absolute average log_2_ fold change of at least 1.

#### Per-cell codon demand analysis

The gene expression (RNA) and valid probe-pair tRNA (tRNA_pp_valid) assays were each log-normalized, reduced to variable features, scaled and subjected to principal component analysis independently, yielding two reductions (pca and tpca). We then constructed a weighted-nearest-neighbor (WNN) graph across both modalities using *FindMultiModalNeighbors*, based on the first 30 principal components of the mRNA reduction and the first 10 of the tRNA reduction. We computed a joint UMAP embedding from the resulting weighted nearest-neighbor graph.

We quantified codon demand per cell by projecting gene expression onto per-gene codon composition. A reference matrix of codon counts per human gene (hg38) was filtered based on the genes present in the gene expression assay, and per-cell codon demand was computed as the matrix product of the transposed codon-count matrix with the log-normalized expression matrix, resulting in a codon-by-cell matrix in which each entry represents the expression-weighted sum of occurrences of that codon across genes. This matrix was added to the Seurat object as an additional assay (“codondemand”) for downstream comparison with the measured tRNA supply.

#### Anticodon representation analysis

We analyzed anticodon-level tRNA counts (assay tRNA_ac) for the unoptimized and betaine-including cell line mixing experiments. For each dataset, the anticodon count matrix (anticodons x cells) was extracted from the Seurat object and pseudobulked by dividing the counts by the total number of cells in the respective dataset, followed by summing of the values across cells. We annotated each anticodon as encoded or non-encoded based on whether a corresponding tRNA gene is annotated in GtRNAdb. We ranked anticodons by mean count per cell in the betaine dataset and assigned the same ranks to features from the standard dataset. For the 13 non-encoded anticodons, we converted mean counts per cell to counts per million within each condition. For each anticodon, we plotted log_10_ CPM values.

#### Quantification of tRNA exon/intron ratios

To estimate how much of each intron-containing tRNA is present in either spliced or unspliced isoform, we designed an additional LHS probe for splice substrates targeting either the intron itself (which is located close to the anticodon within the range of the LHS probe, LHS_intron_) or the splice junction (LHS_exon_), while keeping the joined RHS probe (RHS_common_), resulting in eight exon-intron groups. We compared counts of tRNA features derived from probes targeting either the intron or exon form (ratio LHS_intron_/LHS_exon_).

We split probe pair-level counts from unoptimized and betaine-containing 4-cell-line datasets by cell line. Within each cell line, counts were pseudobulked and for each intron/exon tRNA pair, the intron-to-exon ratio was calculated as the total counts of all intron probe pairs mapping to the same RHS_common_ probe were divided by the counts of the matched exon probe pair. We plotted ratios on a log_10_ scale for each group and cell line.

#### Single-cell guide assignment

gRNA UMI counts were quantified from the kallisto|bustools output (barcodes, genes, and matrix files), imported as sparse matrix, and filtered for cell barcodes that had been called as cells in the matched whole transcriptome library by cellranger processing. We counted each cell’s gRNA UMIs and rated a cell as assignable if exactly one gRNA had 4 or more UMIs, or if one gRNA was detected with at least 2 UMIs and made up at least 75% of the cell’s total UMIs. The Seurat object was filtered for cells passing these criteria and cells were assigned their highest-detected gRNA. UMI-level data were added as assay “sgRNA” to the Seurat object followed by normalization and scaling, and all individual assigned guide names were added to the metadata as the column “identity”, or with de-identified NTC guides (“identity_NTC”).

To validate the efficiency and specificity of our assay, we compared probe-level tRNA expression between perturbed cells and non-targeting controls. Each gRNA was mapped to its target tRNA gene and data were filtered for gene-specific probes, whereby only probe pairs with unique mapping to a tRNA gene were selected. For each gRNA, we compared cells carrying the respective guide to NTC cells using a Wilcoxon rank-sum test on probe-level tRNA expression data with the Seurat function *FindMarkers* (logfc.threshold = 0, min.pct = 0). Results for each target were combined across guides. We plotted the log_2_ fold change either for each guide individually or averaged across guides for each target tRNA gene. The data was plotted as a heatmap of log_2_ fold changes versus NTC (color scale capped at +/- 2.5), filtered for pairs at FDR < 0.05.

#### Isoacceptor and -decoder compensation analysis for tRNA knockdown

For Isoacceptor tRNAs, tRNA composition was calculated within each amino acid family using anticodon-level counts. Each anticodon was mapped to its cognate amino acid and for each cell, the count of the respective anticodon was divided by the summed counts of all anticodons belonging to the same amino acid, transformed to a percentage. We considered a perturbation “targeted” for anticodons that matched the anticodon of the gRNA-targeted isodecoder. The effect was labelled as “isoacceptor” for an anticodon when the gRNA target belonged to the same amino acid group but didn’t match that targeted anticodon and “other” in case of mismatching amino acid identities of the measured tRNA and the gRNA target. Values were plotted as boxplot.

For tRNA-SeC, we measured isotype composition using tRNA counts aggregated by amino acid. For each cell, we divided each amino acid count by the cell’s total tRNA count, generating a per-cell percentage for each amino acid in the total tRNA pool. The data was filtered for cells with guides targeting tRNA-SeC, and for each set of tRNAs grouped by amino acid, the mean % of all tRNA counts per cell were plotted as box plot, with the SeC tRNA group marked in red.

#### Pairwise correlation heatmaps of perturbation profiles

To compare signal responses of the transcriptome and the tRNAome between different guides, we built perturbation-perturbation similarity matrices from pseudobulk mRNA and tRNA profiles. For mRNA, the 2,000 most variable genes were selected from the “RNA” assay using the variance-stabilizing transformation (vst) method in Seurat. We grouped cells by perturbation target, pooling both gRNAs directed against the same target. For each pair, we calculated the pseudobulk profile as the mean UMI count per cell for each variable gene. We then z-scored each gene across all perturbation groups. We calculated similarity between perturbations as the Pearson correlation of their z-scored profiles across genes. We displayed correlation matrices as heatmaps, with rows and columns ordered by hierarchical clustering.

The analysis for the tRNA assay “tRNA_pp_valid” followed the same steps as the transcriptome, except for the variable feature count of 150, adapted to the smaller feature set. To compare the two modalities directly, we drew the tRNA similarity heatmap with rows and columns preserving the order of the mRNA clustering.

#### Primary CD34 differentiation analyses

Starting with the CellRanger filtered feature-barcode matrix, cells were filtered by total counts (min = 1000), genes (min = 500) and mitochondrial content (max = 10%). Genes were retained if detected in ≥55 cells. Raw counts were normalized to the median library size and log-transformed with base 2 and a pseudocount of 1. We selected 4,000 highly variable genes (HVGs) using the highly_variable_genes function in Scanpy (*72*, *73*). We performed principal component analysis (PCA) of log-normalized counts and retained 30 PCs. We clustered cells using PhenoGraph (*74*) through Scanpy with the Leiden algorithm (k = 200, resolution = 1). We then identified cell doublets using Scrublet (*75*) through Scanpy (expected doublet rate = 0.06). We removed any barcodes predicted to be doublets and confirmed that all clusters included fewer than 10% predicted doublets. We then re-processed the filtered matrix using the same steps as above (HVG selection, PCA). We constructed a k-nearest neighbor graph (k = 30) using the top 30 PCs and Euclidean distance. A force-directed layout (FDL) was then computed from the kNN graph using the draw_graph function in Scanpy with default ‘fa’ layout (*76*). We used FDL to visualize cell differentiation because it reflects a continuous manifold rather than discrete cell clusters. We then inferred pseudotime and differentiation trajectories using Palantir (*77*). We selected the top diffusion components (n = 4 based on the eigengap) and assigned a start cell based on high HLF expression (*78*). We defined 3 terminal states as MK, Monocyte, and Erythroid based on expression of marker genes *ITGA2B*, *CD14*, and *GATA1*, respectively (*78*, *79*). We ran Palantir with these manually assigned endpoints and 500 waypoints.

To compare how the transcriptome and tRNAome change along differentiation, we ordered cells within each lineage by Palantir pseudotime and compared the two modalities in their respective principal component spaces. We computed variable features and principal components separately for the gene expression assay (2,000 variable features, 30 PCs) and the valid probe-pair tRNA assay (100 variable features, 30 PCs). Each embedding was centred and each principal component divided by its standard deviation, so that Euclidean distance in the resulting space is equivalent to Mahalanobis distance in the original PC space and a distance of one corresponds to one standard deviation of dataset-wide variation. We used the first 30 PCs for the transcriptome and the first 15 for the tRNAome.

We ranked cells by pseudotime and partitioned them into 100 equally sized bins. We defined the 5% of cells with the lowest and the 5% with the highest pseudotime as the early and late anchor populations. For each bin, we computed the Euclidean distance between the bin centroid and the early anchor centroid, and between the bin centroid and the late anchor centroid, separately in the transcriptome and tRNAome spaces. We obtained standard errors by resampling cells within each bin with replacement (100 bootstrap replicates, fixed seeds), recomputing each distance, and taking the standard deviation across replicates.

For each modality, we calculated the difference between the distance to the early anchor and the distance to the late anchor per bin, and estimated the standard error of the difference as the square root of the sum of the two variances (a conservative estimate that ignores their positive covariance). We smoothed the resulting profiles and their standard errors with a running median (window = 51 bins) and min–max normalized them to the interval [0, 1] independently for each modality. We varied the number of bins and window size to ensure our core conclusion of coordinated changes was robust to hyperparameter selection, and the final parameters reflect a representative demonstration. Shaded bands denote 95% intervals (±1.96 SE). As a negative control, we repeated the analysis with pseudotime values permuted across cells.

#### Trimodal modeling of mRNA, tRNA, and protein data

We processed gene expression, valid probe-pair tRNA measurements, and protein measurements from the same cells independently before integration. Gene expression and tRNA counts were log-normalized, reduced to variable features, scaled and subjected to principal component analysis. Protein counts were normalized by centred log-ratio transformation across features within each cell (*NormalizeData*, normalization.method = “CLR”, margin = 2), scaled and likewise reduced by PCA without variable feature selection. The three reductions were designated pca_mRNA, pca_tRNA and pca_protein.

We then constructed a weighted nearest-neighbor graph across all three modalities with the *FindMultiModalNeighbors* function, using the first 30 principal components of the mRNA reduction and the first 15 of each of the tRNA and protein reductions. We computed a joint UMAP embedding from the weighted nearest-neighbor graph, then clustered cells on the resulting weighted shared-nearest-neighbor graph (wsnn) using the SLM algorithm at a resolution of 0.8.

To test whether tRNA anticodon abundance adds information about protein levels beyond mRNA counts, we fit regression models for all 246 genes with matched protein and RNA levels. We randomly split cells into training (80%) and test (20%) sets. First, normalized protein abundance was corrected for covariates (experimental condition, inferred cell cycle phase, and per-cell protein, mRNA and tRNA read counts) using z-scored residuals from linear regression. We then predicted corrected protein abundance from the gene’s mRNA abundance alone (RNA-only model) versus mRNA abundance plus the cell’s anticodon tRNA pool (RNA+tRNA model). We fit a linear model for the RNA-only model. The RNA+tRNA model used lasso regression for variable selection among the large number predictors. We assessed predictive accuracy using the Pearson correlation between predicted and measured protein abundance in test cells. We defined tRNA-specific information as the difference in those correlations (RNA+tRNA minus RNA-only).

To assess the correlation between a cell’s overall tRNA content and per-cell protein levels, we calculated the tRNA/mRNA ratio for each cell. We selected eight proteins as examples based on the most extreme features on both axes from the combined least squares model For each protein, cells were filtered to non-0 counts for the respective marker in both mRNA and protein assays. A linear regression of protein abundance and mRNA abundance was fitted across these cells, resulting in per-feature residuals. We split cells into quartiles by tRNA:mRNA ratio and plotted them against the residuals.

#### Complex perturbation screening

For the transcriptome-derived joint embedding of each guide target and treatment condition, we pseudobulked raw UMI counts and CPM- and log-transformed them. We ranked genes by variance across all target × condition pseudobulks, selected the top 500 features, and scaled them before principal component analysis. We ran UMAP on the first 30 principal components, selecting UMAP1 and 2 for clustering visualization. The tRNA-based joint embedding of the target x condition data followed the same workflow, except for selecting the top 30 probe pair-level tRNAs for feature selection and 15 principal components for UMAP.

Probe pair-level tRNA counts were summed to pseudobulks for each NTC guide and condition, keeping only pseudobulks of 100 cells or more in each guide/condition combination. We removed probe features with low counts using the edgeR function *filterByExpr*, with treatment as the grouping factor, leaving 216 (AA depletion) and 201 (Torin-1) features. Library sizes were then recalculated and TMM normalization factors applied. Differential expression of tRNA features was tested with limma-voom using sample quality weights (via the voom function *WithQualityWeights*). For each tRNA probe-level feature, we fit a linear model to the voom-transformed log_2_-CPM values with NTC guide and treatment as covariates (∼ guide + treatment), with the control condition as the reference level. We calculated t-statistics using limma’s empirical Bayes procedure (*eBayes*), and adjusted p-values using the Benjamini-Hochberg method. This procedure was applied for both treatment comparisons (AAd/Control and Torin-1/Control), visualized as a correlation plot, and joined DEGs were exported for following analyses.

#### Motif enrichment analysis

Sequences for all high-confidence tRNA genes were extracted from GtRNAdb and the internal promoter elements A and B box were annotated according to previously reported promoter positions (*80*)(**Table S1)**. The full set of 432 tRNA genes was used as input for the function *ggseqlogo* from the R package ggseqlogo to reconstruct the canonical promoter motifs.

For stress-responsiveness enrichment motifs, we extracted differentially regulated tRNAs from the limma-voom analysis described above. The foreground included tRNA probes significantly upregulated (FDR < 0.05, log_2_FC > 0, n = 69) under both AAdepletion and Torin-1 treatment; probes capturing more than one individual promoter sequence were excluded, resulting in 42 filtered probes. The background set included all other retained probes. We assessed nucleotide counts at each A box position for both sets. We added a pseudocount of 0.5 to each count and calculated frequencies. Enrichment was expressed as Kullback-Leibler (KL) divergence for each base, assessing each base’s contribution to the relative entropy of the foreground compared to the background, whereby *p* is the frequency of base *b* at position *i*:

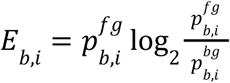

We tested statistical significance for each base and position with a two-sided Fisher’s exact test comparing a given base to the other three possible bases in the foreground vs. background. We adjusted p-values using the Benjamini-Hochberg method. We plotted enrichment values as a sequence logo using the function *ggseglogo*.

## SUPPLEMENTAL TABLES

**TABLE S1.** List of gtRNAdb tRNA gene sequences, per-gene tRNA-targeting LHS and RHS probes, probe pair mapping, and internal promoter sequences.

**TABLE S2.** List of tRNA-gene targeting sgRNA constructs.

**TABLE S3.** List of mRNA-gene targeting sgRNA constructs.

**TABLE S4**. List of CITE-seq antibodies and oligo-barcodes.

**TABLE S5**. Mapping of Amino acid identities to codon logic in DNA, mRNA, and tRNA language.

## CODE AVAILABILITY

Custom code and intermediate data files to reproduce all analyses supporting this manuscript are available at https://github.com/clareaulab/sctrna-reproducibility.

## DATA AVAILABILITY

Sequencing data associated with this work is available at GEO accession **GSE351552** with reviewer access token **cfglwkswlnerjgd**.

## Acknowledgements

We thank members of the Norman, Lareau, and Single-cell Analytics Innovation labs (SAIL) for their helpful feedback. We acknowledge computing assistance from theHigh Performance Computing group and the Integrated Genomics Operation (IGO) Core at Memorial Sloan Kettering Cancer Center. We thank Jason Huang and Kit Nazor of Proteintech Genomics for assistance with the CITE-seq experiments.

## Author contributions

L.C.K., R.C., L.S.L. and C.A.L. conceived and designed the experiments. L.C.K., B.N.N., Y.T.C., A.J.V., K.K., M.M., C.S. and N.M. performed experiments. K.S., E.M., R.C.A., M.T., S.A.V., T.M.N. and L.S.L. provided technical guidance and support for protocol development and optimization. H.V.G. and S.D. provided tissue samples and interpreted data. L.C.K., S.B.B. and A.V. developed software for probe design and data parsing with input from D.P. and C.A.L.; L.C.K. led analyses with feedback from K.K., A.V., D.P., R.C., M.P. and C.A.L. with input from the other authors. L.C.K. and C.A.L. drafted the manuscript with input from all other authors.

## REFERENCES

1. F. Crick, Central dogma of molecular biology. Nature 227, 561–563 (1970).

2. A. Rich, U. L. RajBhandary, Transfer RNA: molecular structure, sequence, and properties. Annu. Rev. Biochem. 45, 805–860 (1976).

3. A. Biela, A. Hammermeister, I. Kaczmarczyk, M. Walczak, L. Koziej, T.-Y. Lin, S. Glatt, The diverse structural modes of tRNA binding and recognition. J. Biol. Chem. 299, 104966 (2023).

4. P. P. Chan, T. M. Lowe, GtRNAdb 2.0: an expanded database of transfer RNA genes identified in complete and draft genomes. Nucleic Acids Res. 44, D184–9 (2016).

5. T. Pan, Modifications and functional genomics of human transfer RNA. Cell Res. 28, 395–404 (2018).

6. A. F. Palazzo, E. S. Lee, Non-coding RNA: what is functional and what is junk? Front. Genet. 6, 2 (2015).

7. P. Thandapani, A. Kloetgen, M. T. Witkowski, C. Glytsou, A. K. Lee, E. Wang, J. Wang, S. E. LeBoeuf, K. Avrampou, T. Papagiannakopoulos, A. Tsirigos, I. Aifantis, Valine tRNA levels and availability regulate complex I assembly in leukaemia. Nature 601, 428–433 (2022).

8. H. Goodarzi, X. Liu, H. C. B. Nguyen, S. Zhang, L. Fish, S. F. Tavazoie, Endogenous tRNA-derived fragments suppress breast cancer progression via YBX1 displacement. Cell 161, 790–802 (2015).

9. S. Chen, D. Markett, M. Karimzadeh, Y. Luo, M. Khoroshkin, B. Boyraz, S. Lee, C. Carpenter, P. Nguyen, K. Garcia, T. Joshi, C. Martin, B. Hänisch, H. Molina, S. Tavazoie, V. Ramani, A. Navickas, H. Goodarzi, Discovery of a tRNA-regulatory transcription factor that suppresses breast cancer metastasis, Genomics (2025). https://www.biorxiv.org/content/10.1101/2025.04.26.650725v1.full.

10. Y. S. Kim, S. Arora, D. Young, A. Tsou, A. Shiuan, C. L. Wladyka, D. Rudoy, J. Y. Kim, J. A. Waters, S. L. Schuster, I. M. Coleman, M. Kapur, M. Sobczyk, C. D. Katanski, A. M. Bayat Tork, W. Zhang, P. S. Nelson, G. Ha, M. C. Haffner, E. Corey, C. Morrissey, Y. Wang, A. R. Subramaniam, J. K. Lee, S. L. Ackerman, T. Pan, A. C. Hsieh, tRNA dosage regulates lineage dependency and resistance in prostate cancer. Nature, doi: 10.1038/s41586-026-11153-8 (2026).

11. J. B. Plotkin, G. Kudla, Synonymous but not the same: the causes and consequences of codon bias. Nat. Rev. Genet. 12, 32–42 (2011).

12. R. G. Roeder, 50+ years of eukaryotic transcription: an expanding universe of factors and mechanisms. Nat Struct Mol Biol 26, 783–791 (2019).

13. I. M. Willis, R. D. Moir, Signaling to and from the RNA Polymerase III Transcription and Processing Machinery. Annu Rev Biochem 87, 75–100 (2018).

14. Z. Bai, D. Zhang, Y. Gao, B. Tao, D. Zhang, S. Bao, A. Enninful, Y. Wang, H. Li, G. Su, X. Tian, N. Zhang, Y. Xiao, Y. Liu, M. Gerstein, M. Li, Y. Xing, J. Lu, M. L. Xu, R. Fan, Spatially exploring RNA biology in archival formalin-fixed paraffin-embedded tissues. Cell 187, 6760–6779.e24 (2024).

15. F. Salmen, J. De Jonghe, T. S. Kaminski, A. Alemany, G. E. Parada, J. Verity-Legg, A. Yanagida, T. N. Kohler, N. Battich, F. van den Brekel, A. L. Ellermann, A. M. Arias, J. Nichols, M. Hemberg, F. Hollfelder, A. van Oudenaarden, High-throughput total RNA sequencing in single cells using VASA-seq. Nat. Biotechnol. 40, 1780–1793 (2022).

16. A. Isakova, D. D. Liu, I. Cvijović, R. Sinha, A. E. Eastman, S. Saul, A. M. Detweiler, N. Neff, S. Einav, I. L. Weissman, S. R. Quake, Scalable single-cell total RNA sequencing unifies coding and noncoding transcriptomics. Nat. Biotechnol., 1–12 (2026).

17. M. Stoeckius, C. Hafemeister, W. Stephenson, B. Houck-Loomis, P. K. Chattopadhyay, H. Swerdlow, R. Satija, P. Smibert, Simultaneous epitope and transcriptome measurement in single cells. Nat. Methods 14, 865–868 (2017).

18. M. Külp, M. Osuna Lopez, A.-S. Wiegand, S. P. Perner, R. Cetin, Y. Matthess, S. Günther, T. Schmachtel, H. Schultheis, M. Looso, M. Kaulich, B. Häupl, T. Oellerich, V. Benes, H. Bonig, M. A. Rieger, Single-cell mapping of tRNA expression dynamics across human hematopoiesis, bioRxiv (2026). 10.64898/2026.08.14.744819.

19. A. Dixit, O. Parnas, B. Li, J. Chen, C. P. Fulco, L. Jerby-Arnon, N. D. Marjanovic, D. Dionne, T. Burks, R. Raychowdhury, B. Adamson, T. M. Norman, E. S. Lander, J. S. Weissman, N. Friedman, A. Regev, Perturb-seq: Dissecting molecular circuits with scalable single-cell RNA profiling of pooled genetic screens. Cell 167, 1853–1866.e17 (2016).

20. N. H. Padhiar, U. Katneni, A. A. Komar, Y. Motorin, C. Kimchi-Sarfaty, Advances in methods for tRNA sequencing and quantification. Trends Genet 40, 276–290 (2024).

21. S. B. Blattman, N. Maslah, A. A. Varela, K. Kumpaitis, B. Nalbant, C. Snopkowski, M. Mariani, L. C. Kida, M. Takizawa, N. Ratnayeke, K. K. H. Yu, S. Fernandes, N. Mousavi, E. Borgstrom, D. Vallejo, L. Boghospor, R. Xin, M. Mignardi, S. Wu, N. Scarlott, L. Delgado-Rivera, P. Kumar, S. Krishnan, S. Giraudier, J.-J. Kiladjian, B. E. Howitt, A. Kohlway, P. Lund, D. Pe’er, R. Chaligne, C. A. Lareau, Scalable genotyping in fixed transcriptomes resolves clonal heterogeneity via single-cell sequencing, bioRxivorg (2026). 10.64898/2026.04.11.717967.

22. T. Abay, R. R. Stickels, M. T. Takizawa, B. N. Nalbant, Y.-H. Hsieh, S. Hwang, C. Snopkowski, K. K. H. Yu, Z. Abou-Mrad, V. Tabar, B. E. Howitt, L. S. Ludwig, R. Chaligné, A. T. Satpathy, C. A. Lareau, Transcript-specific enrichment enables profiling of rare cell states via single-cell RNA sequencing. Nat. Genet., 1–10 (2025).

23. M. Pak, M. S. Saurty-Seerunghen, K. Wise, T.-A. Abera, C. Lama, N. Parghi, T. Kang, X. Sun, Q. Gao, L. Bao, M. Roshal, J. N. Allan, R. R. Furman, L. G. Martelotto, A. S. Nam, Co-mapping clonal and transcriptional heterogeneity in somatic evolution via GoT-Multi. Cell Genom. 6, 101036 (2026).

24. A. Bradu, J. D. Blair, I. N. Grabski, I. Mascio, J. Lee, C. McCormick, R. Satija, Genome-wide single-cell perturbation screens with VIPerturb-seq, bioRxivorg (2026). 10.64898/2026.02.12.705613.

25. M. F. de Oliveira, J. P. Romero, M. Chung, S. R. Williams, A. D. Gottscho, A. Gupta, S. E. Pilipauskas, S. Mohabbat, N. Raman, D. J. Sukovich, D. M. Patterson, Visium HD Development Team, S. E. B. Taylor, High-definition spatial transcriptomic profiling of immune cell populations in colorectal cancer. Nat. Genet. 57, 1512–1523 (2025).

26. M. Y. Pavlov, R. E. Watts, Z. Tan, V. W. Cornish, M. Ehrenberg, A. C. Forster, Slow peptide bond formation by proline and other N-alkylamino acids in translation. Proceedings of the National Academy of Sciences 106, 50–54 (2009).

27. R. J. Argüello, A. J. Combes, R. Char, J.-P. Gigan, A. I. Baaziz, E. Bousiquot, V. Camosseto, B. Samad, J. Tsui, P. Yan, S. Boissonneau, D. Figarella-Branger, E. Gatti, E. Tabouret, M. F. Krummel, P. Pierre, SCENITH: A flow cytometry-based method to functionally profile energy metabolism with single-cell resolution. Cell Metab. 32, 1063–1075.e7 (2020).

28. R. A. Saunders, W. E. Allen, X. Pan, J. Sandhu, J. Lu, T. K. Lau, K. Smolyar, Z. A. Sullivan, C. Dulac, J. S. Weissman, X. Zhuang, Perturb-Multimodal: A platform for pooled genetic screens with imaging and sequencing in intact mammalian tissue. Cell 188, 4790–4809.e22 (2025).

29. J. D. Blair, A. Bradu, C. Dalgarno, I. N. Grabski, R. Satija, Iterative, multimodal, and scalable single-cell profiling for discovery and characterization of signaling regulators, bioRxivorg (2025). 10.1101/2025.08.27.672635.

30. A. Dixit, O. Parnas, B. Li, J. Chen, C. P. Fulco, L. Jerby-Arnon, N. D. Marjanovic, D. Dionne, T. Burks, R. Raychowdhury, B. Adamson, T. M. Norman, E. S. Lander, J. S. Weissman, N. Friedman, A. Regev, Perturb-Seq: Dissecting Molecular Circuits with Scalable Single-Cell RNA Profiling of Pooled Genetic Screens. Cell 167, 1853–1866.e17 (2016).

31. J. M. Replogle, R. A. Saunders, A. N. Pogson, J. A. Hussmann, A. Lenail, A. Guna, L. Mascibroda, E. J. Wagner, K. Adelman, G. Lithwick-Yanai, N. Iremadze, F. Oberstrass, D. Lipson, J. L. Bonnar, M. Jost, T. M. Norman, J. S. Weissman, Mapping information-rich genotype-phenotype landscapes with genome-scale Perturb-seq. Cell 185, 2559–2575.e28 (2022).

32. Y. Kawatani, T. Suzuki, R. Shimizu, V. P. Kelly, M. Yamamoto, Nrf2 and selenoproteins are essential for maintaining oxidative homeostasis in erythrocytes and protecting against hemolytic anemia. Blood 117, 986–996 (2011).

33. C. Liao, R. C. Hardison, M. J. Kennett, B. A. Carlson, R. F. Paulson, K. S. Prabhu, Selenoproteins regulate stress erythroid progenitors and spleen microenvironment during stress erythropoiesis. Blood 131, 2568–2580 (2018).

34. D. W. Smith, A. L. McNamara, Specialization of rabbit reticulocyte transfer RNA content for hemoglobin synthesis. Science 171, 577–579 (1971).

35. H. Gingold, D. Tehler, N. R. Christoffersen, M. M. Nielsen, F. Asmar, S. M. Kooistra, N. S. Christophersen, L. L. Christensen, M. Borre, K. D. Sørensen, L. D. Andersen, C. L. Andersen, E. Hulleman, T. Wurdinger, E. Ralfkiær, K. Helin, K. Grønbæk, T. Ørntoft, S. M. Waszak, O. Dahan, J. S. Pedersen, A. H. Lund, Y. Pilpel, A dual program for translation regulation in cellular proliferation and differentiation. Cell 158, 1281–1292 (2014).

36. J. D. Buenrostro, M. R. Corces, C. A. Lareau, B. Wu, A. N. Schep, M. J. Aryee, R. Majeti, H. Y. Chang, W. J. Greenleaf, Integrated Single-Cell Analysis Maps the Continuous Regulatory Landscape of Human Hematopoietic Differentiation. Cell 173, 1535–1548.e16 (2018).

37. J. D. Buenrostro, M. R. Corces, C. A. Lareau, B. Wu, A. N. Schep, M. J. Aryee, R. Majeti, H. Y. Chang, W. J. Greenleaf, Integrated single-cell analysis maps the continuous regulatory landscape of human hematopoietic differentiation. Cell 173, 1535–1548.e16 (2018).

38. L. Velten, S. F. Haas, S. Raffel, S. Blaszkiewicz, S. Islam, B. P. Hennig, C. Hirche, C. Lutz, E. C. Buss, D. Nowak, T. Boch, W.-K. Hofmann, A. D. Ho, W. Huber, A. Trumpp, M. A. G. Essers, L. M. Steinmetz, Human haematopoietic stem cell lineage commitment is a continuous process. Nat. Cell Biol. 19, 271–281 (2017).

39. S. C. Bendall, K. L. Davis, E.-A. D. Amir, M. D. Tadmor, E. F. Simonds, T. J. Chen, D. K. Shenfeld, G. P. Nolan, D. Pe’er, Single-cell trajectory detection uncovers progression and regulatory coordination in human B cell development. Cell 157, 714–725 (2014).

40. S. Ma, B. Zhang, L. M. LaFave, A. S. Earl, Z. Chiang, Y. Hu, J. Ding, A. Brack, V. K. Kartha, T. Tay, T. Law, C. Lareau, Y.-C. Hsu, A. Regev, J. D. Buenrostro, Chromatin Potential Identified by Shared Single-Cell Profiling of RNA and Chromatin. Cell 183, 1103–1116.e20 (2020).

41. L. Anderson, J. Seilhamer, A comparison of selected mRNA and protein abundances in human liver. Electrophoresis 18, 533–537 (1997).

42. B. Schwanhäusser, D. Busse, N. Li, G. Dittmar, J. Schuchhardt, J. Wolf, W. Chen, M. Selbach, Global quantification of mammalian gene expression control. Nature 473, 337–342 (2011).

43. T. Maier, M. Güell, L. Serrano, Correlation of mRNA and protein in complex biological samples. FEBS Letters 583, 3966–3973 (2009).

44. K. Asano, J. Clayton, A. Shalev, A. G. Hinnebusch, A multifactor complex of eukaryotic initiation factors, eIF1, eIF2, eIF3, eIF5, and initiator tRNA(Met) is an important translation initiation intermediate in vivo. Genes Dev 14, 2534–2546 (2000).

45. M. Pavon-Eternod, S. Gomes, M. R. Rosner, T. Pan, Overexpression of initiator methionine tRNA leads to global reprogramming of tRNA expression and increased proliferation in human epithelial cells. RNA 19, 461–466 (2013).

46. C. J. Clarke, T. J. Berg, J. Birch, D. Ennis, L. Mitchell, C. Cloix, A. Campbell, D. Sumpton, C. Nixon, K. Campbell, V. L. Bridgeman, P. B. Vermeulen, S. Foo, E. Kostaras, J. L. Jones, L. Haywood, E. Pulleine, H. Yin, D. Strathdee, O. Sansom, K. Blyth, I. McNeish, S. Zanivan, A. R. Reynolds, J. C. Norman, The Initiator Methionine tRNA Drives Secretion of Type II Collagen from Stromal Fibroblasts to Promote Tumor Growth and Angiogenesis. Curr Biol 26, 755–765 (2016).

47. R. K. Shrimali, R. D. Irons, B. A. Carlson, Y. Sano, V. N. Gladyshev, J. M. Park, D. L. Hatfield, Selenoproteins mediate T cell immunity through an antioxidant mechanism. J Biol Chem 283, 20181–20185 (2008).

48. K. A. Spriggs, M. Bushell, A. E. Willis, Translational regulation of gene expression during conditions of cell stress. Mol Cell 40, 228–237 (2010).

49. M. Costa-Mattioli, P. Walter, The integrated stress response: From mechanism to disease. Science 368 (2020).

50. D. Canella, V. Praz, J. H. Reina, P. Cousin, N. Hernandez, Defining the RNA polymerase III transcriptome: Genome-wide localization of the RNA polymerase III transcription machinery in human cells. Genome Res 20, 710–721 (2010).

51. D. Canella, D. Bernasconi, F. Gilardi, G. LeMartelot, E. Migliavacca, V. Praz, P. Cousin, M. Delorenzi, N. Hernandez, CycliX Consortium, A multiplicity of factors contributes to selective RNA polymerase III occupancy of a subset of RNA polymerase III genes in mouse liver. Genome Res 22, 666–680 (2012).

52. R. Ferrari, C. Rivetti, J. Acker, G. Dieci, Distinct roles of transcription factors TFIIIB and TFIIIC in RNA polymerase III transcription reinitiation. Proc Natl Acad Sci U S A 101, 13442–13447 (2004).

53. W. Seifert-Davila, M. Girbig, L. Hauptmann, T. Hoffmann, S. Eustermann, C. W. Müller, Structural insights into human TFIIIC promoter recognition. Sci Adv 9, eadh2019 (2023).

54. R. Ferrari, G. Dieci, The transcription reinitiation properties of RNA polymerase III in the absence of transcription factors. Cell Mol Biol Lett 13, 112–118 (2008).

55. C. Bunne, Y. Roohani, Y. Rosen, A. Gupta, X. Zhang, M. Roed, T. Alexandrov, M. AlQuraishi, P. Brennan, D. B. Burkhardt, A. Califano, J. Cool, A. F. Dernburg, K. Ewing, E. B. Fox, M. Haury, A. E. Herr, E. Horvitz, P. D. Hsu, V. Jain, G. R. Johnson, T. Kalil, D. R. Kelley, S. O. Kelley, A. Kreshuk, T. Mitchison, S. Otte, J. Shendure, N. J. Sofroniew, F. Theis, C. V. Theodoris, S. Upadhyayula, M. Valer, B. Wang, E. Xing, S. Yeung-Levy, M. Zitnik, T. Karaletsos, A. Regev, E. Lundberg, J. Leskovec, S. R. Quake, How to build the virtual cell with artificial intelligence: Priorities and opportunities. Cell 187, 7045–7063 (2024).

56. X. Zhang, G. V. Shivashankar, C. Uhler, Partially shared multi-modal embedding learns holistic representation of cell state. Nat Comput Sci 6, 285–300 (2026).

57. A. J. Ashford, T. Enright, J. Somers, O. Nikolova, E. Demir, Unifying multimodal single-cell data with a mixture-of-experts β-variational autoencoder framework. Genome Res 36, 2068–2090 (2026).

58. F. Salmen, J. De Jonghe, T. S. Kaminski, A. Alemany, G. E. Parada, J. Verity-Legg, A. Yanagida, T. N. Kohler, N. Battich, F. van den Brekel, A. L. Ellermann, A. M. Arias, J. Nichols, M. Hemberg, F. Hollfelder, A. van Oudenaarden, High-throughput total RNA sequencing in single cells using VASA-seq. Nat Biotechnol 40, 1780–1793 (2022).

59. A. Isakova, D. D. Liu, I. Cvijović, R. Sinha, A. E. Eastman, S. Saul, A. M. Detweiler, N. Neff, S. Einav, I. L. Weissman, S. R. Quake, Scalable single-cell total RNA sequencing unifies coding and noncoding transcriptomics. Nat Biotechnol, doi: 10.1038/s41587-026-03068-6 (2026).

60. M. E. Evans, W. C. Clark, G. Zheng, T. Pan, Determination of tRNA aminoacylation levels by high-throughput sequencing. Nucleic Acids Res 45, e133 (2017).

61. A. Behrens, G. Rodschinka, D. D. Nedialkova, High-resolution quantitative profiling of tRNA abundance and modification status in eukaryotes by mim-tRNAseq. Mol Cell 81, 1802–1815.e7 (2021).

62. P. Boccaletto, F. Stefaniak, A. Ray, A. Cappannini, S. Mukherjee, E. Purta, M. Kurkowska, N. Shirvanizadeh, E. Destefanis, P. Groza, G. Avşar, A. Romitelli, P. Pir, E. Dassi, S. G. Conticello, F. Aguilo, J. M. Bujnicki, MODOMICS: a database of RNA modification pathways. 2021 update. Nucleic Acids Res 50, D231–D235 (2022).

63. L. B. Earnest-Noble, D. Hsu, S. Chen, H. Asgharian, M. Nandan, M. C. Passarelli, H. Goodarzi, S. F. Tavazoie, Two isoleucyl tRNAs that decode synonymous codons divergently regulate breast cancer metastatic growth by controlling translation of proliferation-regulating genes. Nat Cancer 3, 1484–1497 (2022).

64. B. Xie, J. Chen, Z. Wang, Q. Yin, Z.-M. Dai, Sweet enhancers of polymerase chain reaction. PLoS One 19, e0311939 (2024).

65. J. G. Doench, N. Fusi, M. Sullender, M. Hegde, E. W. Vaimberg, K. F. Donovan, I. Smith, Z. Tothova, C. Wilen, R. Orchard, H. W. Virgin, J. Listgarten, D. E. Root, Optimized sgRNA design to maximize activity and minimize off-target effects of CRISPR-Cas9. Nat Biotechnol 34, 184–191 (2016).

66. K. M. Southard, R. C. Ardy, A. Tang, D. D. O’Sullivan, E. Metzner, K. Guruvayurappan, T. M. Norman, Comprehensive transcription factor perturbations recapitulate fibroblast transcriptional states. Nat Genet 57, 2323–2334 (2025).

67. A. Tang, R. C. Ardy, R. E. Mendes, T. M. Norman, Scaling perturbations: beyond genome-scale CRISPR screens, bioRxiv (2026). 10.64898/2026.01.16.699948.

68. M. Martin, Cutadapt removes adapter sequences from high-throughput sequencing reads. EMBnet J. 17, 10 (2011).

69. N. L. Bray, H. Pimentel, P. Melsted, L. Pachter, Near-optimal probabilistic RNA-seq quantification. Nat Biotechnol 34, 525–527 (2016).

70. P. Melsted, V. Ntranos, L. Pachter, The barcode, UMI, set format and BUStools. Bioinformatics 35, 4472–4473 (2019).

71. I. Tirosh, B. Izar, S. M. Prakadan, M. H. Wadsworth II, D. Treacy, J. J. Trombetta, A. Rotem, C. Rodman, C. Lian, G. Murphy, M. Fallahi-Sichani, K. Dutton-Regester, J.-R. Lin, O. Cohen, P. Shah, D. Lu, A. S. Genshaft, T. K. Hughes, C. G. K. Ziegler, S. W. Kazer, A. Gaillard, K. E. Kolb, A.-C. Villani, C. M. Johannessen, A. Y. Andreev, E. M. Van Allen, M. Bertagnolli, P. K. Sorger, R. J. Sullivan, K. T. Flaherty, D. T. Frederick, J. Jané-Valbuena, C. H. Yoon, O. Rozenblatt-Rosen, A. K. Shalek, A. Regev, L. A. Garraway, Dissecting the multicellular ecosystem of metastatic melanoma by single-cell RNA-seq. Science, doi: 10.1126/science.aad0501 (2016).

72. F. A. Wolf, P. Angerer, F. J. Theis, SCANPY: large-scale single-cell gene expression data analysis. Genome Biol. 19, 15 (2018).

73. T. Stuart, A. Butler, P. Hoffman, C. Hafemeister, E. Papalexi, W. M. Mauck 3rd, Y. Hao, M. Stoeckius, P. Smibert, R. Satija, Comprehensive integration of single-cell data. Cell 177, 1888–1902.e21 (2019).

74. J. H. Levine, E. F. Simonds, S. C. Bendall, K. L. Davis, E.-A. D. Amir, M. D. Tadmor, O. Litvin, H. G. Fienberg, A. Jager, E. R. Zunder, R. Finck, A. L. Gedman, I. Radtke, J. R. Downing, D. Pe’er, G. P. Nolan, Data-driven phenotypic dissection of AML reveals progenitor-like cells that correlate with prognosis. Cell 162, 184–197 (2015).

75. S. L. Wolock, R. Lopez, A. M. Klein, Scrublet: Computational identification of cell Doublets in Single-cell transcriptomic data. Cell Syst. 8, 281–291.e9 (2019).

76. M. Jacomy, T. Venturini, S. Heymann, M. Bastian, ForceAtlas2, a continuous graph layout algorithm for handy network visualization designed for the Gephi software. PLoS One 9, e98679 (2014).

77. M. Setty, V. Kiseliovas, J. Levine, A. Gayoso, L. Mazutis, D. Pe’er, Characterization of cell fate probabilities in single-cell data with Palantir. Nat. Biotechnol. 37, 451–460 (2019).

78. D. Pellin, M. Loperfido, C. Baricordi, S. L. Wolock, A. Montepeloso, O. K. Weinberg, A. Biffi, A. M. Klein, L. Biasco, A comprehensive single cell transcriptional landscape of human hematopoietic progenitors. Nat. Commun. 10, 2395 (2019).

79. S. Triana, D. Vonficht, L. Jopp-Saile, S. Raffel, R. Lutz, D. Leonce, M. Antes, P. Hernández-Malmierca, D. Ordoñez-Rueda, B. Ramasz, T. Boch, J.-C. Jann, D. Nowak, W.-K. Hofmann, C. Müller-Tidow, D. Hübschmann, T. Alexandrov, V. Benes, A. Trumpp, M. Paulsen, L. Velten, S. Haas, Single-cell proteo-genomic reference maps of the hematopoietic system enable the purification and massive profiling of precisely defined cell states. Nat. Immunol. 22, 1577–1589 (2021).

80. E. A. Savina, T. G. Shumilina, V. A. Porolo, G. S. Lebedev, Y. L. Orlov, A. A. Anashkina, I. A. Il’icheva, Structural Features of DNA in tRNA Genes and Their Upstream Sequences. Int J Mol Sci 25 (2024).

